# Promoter mutagenesis and a massively parallel reporter screen of the *MAPT* locus identifies cis-regulatory elements and genetic variation effects

**DOI:** 10.64898/2026.03.06.710116

**Authors:** Rebecca M. Hauser, Henry L. Limbo, J. Nicholas Brazell, Belle A. Moyers, Shelby N. Lauzon, Erin A. Barinaga, S. Quinn Johnston, Brianne B. Rogers, Jared W. Taylor, J. Nicholas Cochran

## Abstract

Tau neurofibrillary tangles are a hallmark of several neurodegenerative diseases called tauopathies, including frontotemporal dementia and Alzheimer’s Disease. Ongoing clinical trials for tauopathies seek to reduce Tau in the brain through immunotherapy, antisense oligonucleotides, and siRNA. *MAPT* codes for Tau, therefore understanding how the *MAPT* gene is regulated and the effect of genetic variation at its regulatory elements is likely to have high relevance for tauopathies. We screened a ∼3 Mb region including the *MAPT* locus using 2 different massively parallel reporter assay (MPRA) strategies in KOLF2.1J h-NGN2 neurons and HEK293FT cells, identifying previously unannotated cis-regulatory elements (CREs). Using CRISPR interference (CRISPRi) in mixed neuron cultures, we identified a new CRE for *MAPT*, as well as 2 CREs for another nearby gene of interest, *KANSL1*. Known genetic variation from the Alzheimer’s Disease sequencing project was tested in a separate MPRA at the top CREs near the *MAPT* gene, identifying variants with altered regulatory effects including those at previously identified CREs for *MAPT*. Using a saturation mutagenesis screen of a 2,000 bp region encompassing the *MAPT* promoter, we assessed regulatory effects of each possible single nucleotide variant in this region. We identified several neuron-specific regulatory variant effects at this region, including a high confidence binding site for the transcription factors EGR2, ZBTB14, and TCLF5 at a region of high MPRA activity and genetic conservation.

## Introduction

Tau is a microtubule associated protein encoded by the gene *MAPT* whose dysregulation plays an important role in several neurodegenerative disorders called tauopathies, including progressive supranuclear palsy (PSP), frontotemporal dementia (FTD), and Alzheimer’s Disease (AD). Alzheimer’s disease is the most common tauopathy, and is considered a secondary tauopathy with the primary pathology being amyloid beta plaques.^1^ In AD, Tau becomes hyperphosphorylated leading to the formation of aggregates called neurofibrillary tangles that precede cell death.^2^ Over time, the burden of Tau aggregates and amyloid plaques increases and correlates with neurodegeneration and the impairment of cognitive ability.

Recently, reduction of Tau has gained interest as a potential target for tauopathy therapeutics. Tau reduction has been shown to have a beneficial effect in animal models of AD without severe deficits.^3–8^ Several pharmaceutical companies have tested immunotherapies targeting Tau with the goal of clearing the pathological protein from the brain.^9–20^ Another strategy being explored to reduce total Tau is targeting *MAPT* with antisense oligonucleotide or siRNA therapy to reduce overall gene expression level.^21–23^ To date, 15 therapies targeting Tau pathology are in development.^24,25^ Due to its shared pathology across multiple diseases and advances in technology allowing us to develop more targeted reduction strategies, it has become increasingly important to understand how *MAPT* is regulated.

The *MAPT* gene is located on chromosome 17 within a well known chromosomal inversion region at 17q21.31. This inversion is sorted into two primary haplotypes H1 (forward orientation) and H2 (inverted orientation). The H1 haplotype is associated with an increased risk for neurodegenerative disorders including Alzheimer’s disease, while the H2 haplotype is considered to have a protective effect in AD.^26^ The primary *MAPT* promoter is GC rich without a TATA box domain, and is located upstream of the first exon of the *MAPT* gene encoding for the 5’UTR.^27,28^ Tau has several different isoforms with unique properties. Alternate splicing of exon 10 results in either 3R or 4R Tau named for the number of repeat domains in the microtubule binding region.^1^ Different tauopathies have varying ratios of 3R and 4R Tau with Alzheimer’s disease typically having a ratio of 1:1 3R to 4R Tau, and progressive supranuclear palsy showing primarily 4R Tau pathology.^29^ In addition, inclusion of exon 4a results in an isoform known as “big Tau.”^30^ Big Tau is primarily located in peripheral neurons^31–33^, and it is possible that its larger size could promote axonal transport. Furthermore, this isoform might be protective against forming aggregates due to a reduction in phosphorylation sites.^34–38^

Recently, we identified several cis-regulatory elements (CREs) for *MAPT.*^39^ We sought to expand our initial study to further understand *MAPT* regulation by performing an unbiased screen of the *MAPT* locus to identify regulatory elements using massively parallel reporter assays (MPRAs). Additionally, we evaluated the effects of genetic variation at the *MAPT* promoter through saturation mutagenesis of a 2,000 bp region encompassing the promoter and evaluated known genetic variation from the Alzheimer’s Disease Sequencing Project (ADSP) database at validated and proximal CREs for *MAPT*. This work increases our understanding of Tau regulation and adds to the growing resource of functional regulatory element screens in neurons.^40–48^

## Results

### MPRA screens identify new cell-type specific CREs

To screen the *MAPT* locus for cis-regulatory elements, we utilized two massively parallel reporter assay strategies. For both MPRAs, the lentiMPRA plasmid was transfected into HEK293FT cells (HEKs) to generate lentivirus to transduce 14 day old KOLF 2.1j-hNGN2 neurons (fig. 1a.).^49,50^ Bacterial artificial chromosomes (BACs) aligning to a ∼3 Mb region encompassing the *MAPT* locus were selected and randomly sheared to an average size of 250 bp to use as input for an MPRA (BAC MPRA). A synthesized oligo pool strategy (Oligo MPRA) was implemented for regions of low coverage from the BAC MPRA, including a region directly covering the *MAPT* gene as well as for other regions of interest. 270 bp oligos were designed every 90 bp tiling the forward and reverse strands of genomic DNA in this region and ordered from Twist Biosciences (fig. 1b). For both MPRAs, we observed that the majority of CREs were specific to each tested cell type. We identified 1,636 bins with positive regulatory activity in neurons and 2,257 in HEKs with an overlap of 266 shared bins per cell type in the BAC MPRA (pval.mad ≤ 0.05) (fig. 1c, tables s2-3). We identified 2,900 oligos with significant positive regulatory activity in neurons and 328 oligos in HEKs with an overlap of 84 shared between the two cell types in the Oligo MPRA (pval.mad ≤ 0.05) (fig. 1d, tables s4-5). When overlapping and consecutive significant bins/oligos were joined to form a single CRE, we identified 1,335 CREs using the BAC MPRA in neurons and 1,402 CREs in the HEKs as well as 864 CREs in the Oligo MPRA in neurons and 254 in HEKs (fig. s1). Comparing our MPRA data to annotated ENCODE v4 cCREs^51^, the majority of CREs found using the BAC MPRA were previously unannotated (fig. 1e), and the majority of the CREs found using the Oligo MPRA were annotated as enhancer-like sequences (dELS) (fig. 1f).

**Figure 1.**
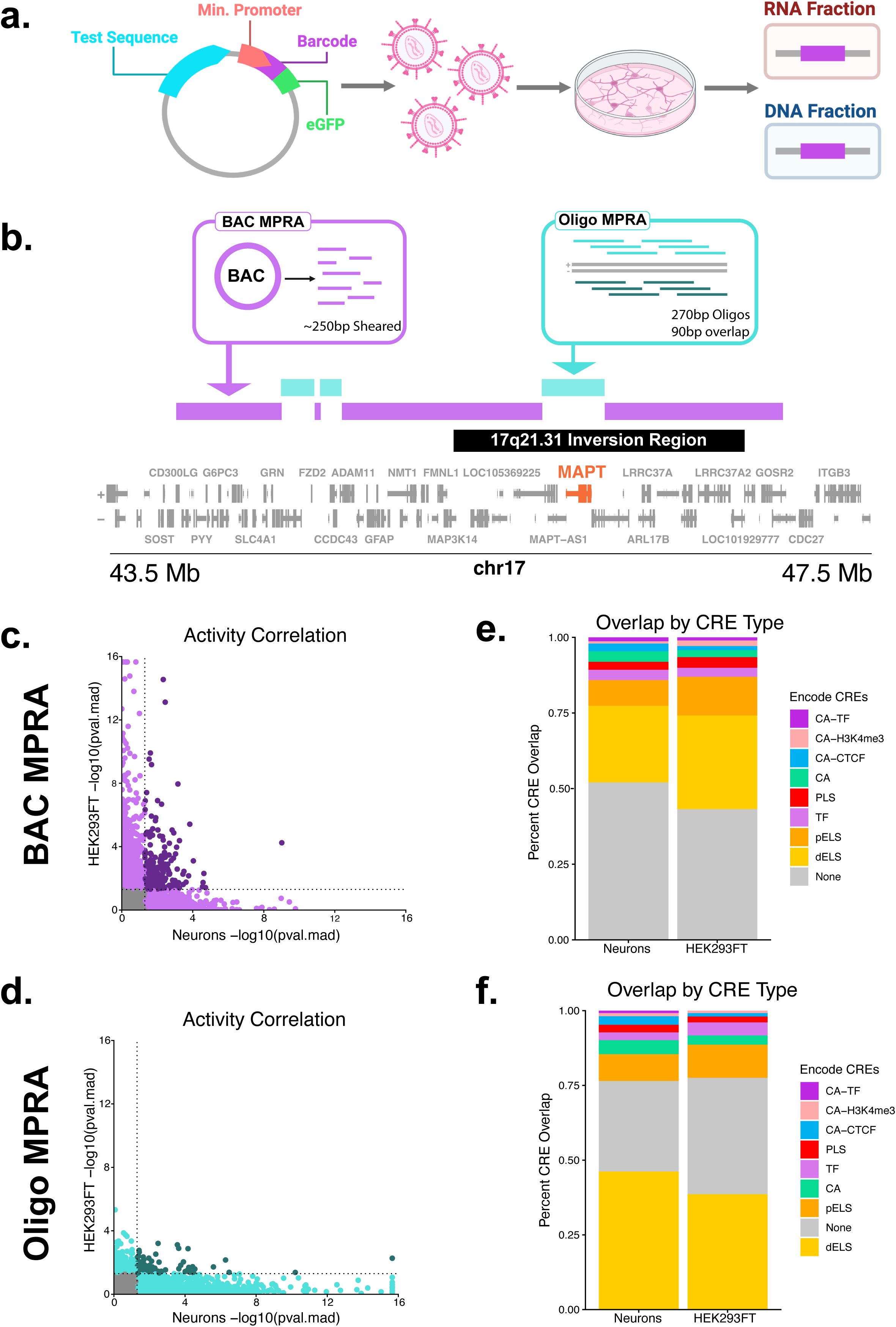
**a.** LentiMPRA experimental design diagram. **b.** Gene track of the *MAPT* locus and regions of the genome covered by the BAC and Oligo MPRAs. Gaps in the BAC MPRA represent regions of low MPRA coverage where the data was unusable. The Oligo MPRA was designed to supplement these gaps. **c.** Correlation of regulatory activity of 100 bp genomic bins from the BAC MPRA in Neurons and HEK293FT cells. (mad.score Spearman’s ⍴ = 0.0162, approximate p = 0.0395). Pval.mad was capped at a lower limit of 2.2×10^-16^. **d.** Correlation of regulatory activity of 270 bp oligos from the Oligo MPRA in Neurons and HEK293FT cells. (mad.score Spearman’s ⍴ = -0.0464, approximate p = 2.86×10^-4^). **e.** Overlap of ENCODE v4 cCREs with regions identified as regulatory elements through the BAC MPRA (pval.mad ≤ 0.05). Pval.mad was capped at a lower limit of 2.2×10^-16^**. f.** Overlap of ENCODE v4 cCREs with regions identified as regulatory elements through the Oligo MPRA (pval.mad ≤ 0.05). CA-TF = chromatin accessibility with transcription factor binding, CA-H3K4me3 = chromatin accessibility with H3K4me3, CA-CTF = chromatin accessibility with CTCF, CA = chromatin accessibility, PLS = promoter-like sequence, TF = transcription factor binding, pELS = proximal enhancer-like sequence, dELS = distal enhancer-like sequence.

### CRISPRi identifies new CREs for MAPT and KANSL1

From the CREs identified in the initial MPRA screens, we nominated 14 regions as potential regulatory elements for *MAPT* by considering overlap of previously published 3D -omics data^39^ and multiomics data^39,52^ and incorporating both the distance from the *MAPT* promoter and the strength of the enhancer activity in our MPRAs (fig. 2a).^39,52^ CRISPRi guides were designed to each region and were transduced into KOLF 2.1J NPCs.^39^ NPCs were differentiated following the the Bardy et al. protocol,^53^ which produces a mixed culture of inhibitory and excitatory neurons, as well as some immature astrocytes. RNA was collected from the cells and sent for mRNA-sequencing. Because *MAPT* is expressed in both neurons and astrocytes, to account for the multiple cell types present in the bulk RNA as well as well-to-well cell count variability, Seurat^54^ cell type scores for neurons and astrocytes were added to the DESeq2^55^ model to ensure that changes in *MAPT* expression were due to *MAPT* repression and not a reduction in total neuron count (table s9). Expression of genes within 200 kb of the guides were examined to look for likely gene targets by proximity of the candidate enhancers (fig. 2b).^56^ We discovered a novel enhancer for *MAPT* at region r5 (adj. p. value = 2.24×10^-4^ and 0.0213 before and after accounting for cell type score) ∼113,349 bp upstream of the *MAPT* TSS falling within a *MAPT* promoter Capture-C link identified in cultured neurons.^39^ Excluding the cell type score from the DESeq2 model, repression of region r8 located ∼72,708 bp upstream of the *MAPT* TSS within a *MAPT* promoter HiC link also resulted in a significant reduction of *MAPT* expression (adj. p. value = 0.0145 and 0.990 before and after accounting for cell type score). This is likely due to an overall reduction of neuron count caused by a prevention of differentiation of NPCs into neurons following the blocking of this CRE with CRISPRi. We are reporting this region as a CRE important for neuronal differentiation and can not exclude that it is acting directly on *MAPT* regulation (fig. 2c). In the process of testing these CREs, we found regions r12 and r13 to be CREs for the gene *KANSL1* (r12 adj. p. values = 4.86×10^-6^, 2.94×10^-6^ before and after adjusting for cell type score, and r13 adj. p. values < 2.2×10^-16^ both before and after adjusting for cell type score). Pathogenic variants within *KANSL1* are known to cause the developmental disorder Koolen-de Vries Syndrome.^57^ In addition, variants at the *KANSL1* locus in connection with H1/H2 haplotypes are associated through genome-wide association studies with increased AD, Parkinson’s disease, and amyotrophic lateral sclerosis risk.^58,59^ Region r5, in addition to being a high confidence CRE for *MAPT* also decreased *KANSL1* expression when targeted by CRISPRi, although this association was lost once accounting for cell type score (fig. 2d) (adj. p. value = 2.03 x 10^-3^ and 0.118 before and after adjusting for cell type score). Several CREs affected expression of the *GFAP* gene. We believe that region r1 (adj. p. value = 3.68×10^-12^ and 1.07×10^-16^ before and after adjusting for cell type score) could be a CRE for GFAP due to its proximity to the *GFAP* promoter (between 2 ENCODE CREs within *GFAP* exon 9^60^), however we cannot be certain if the other CREs are directly influencing *GFAP* expression or indirectly affecting it through other mechanisms including altering astrocyte reactivity.^61^

**Figure 2.**
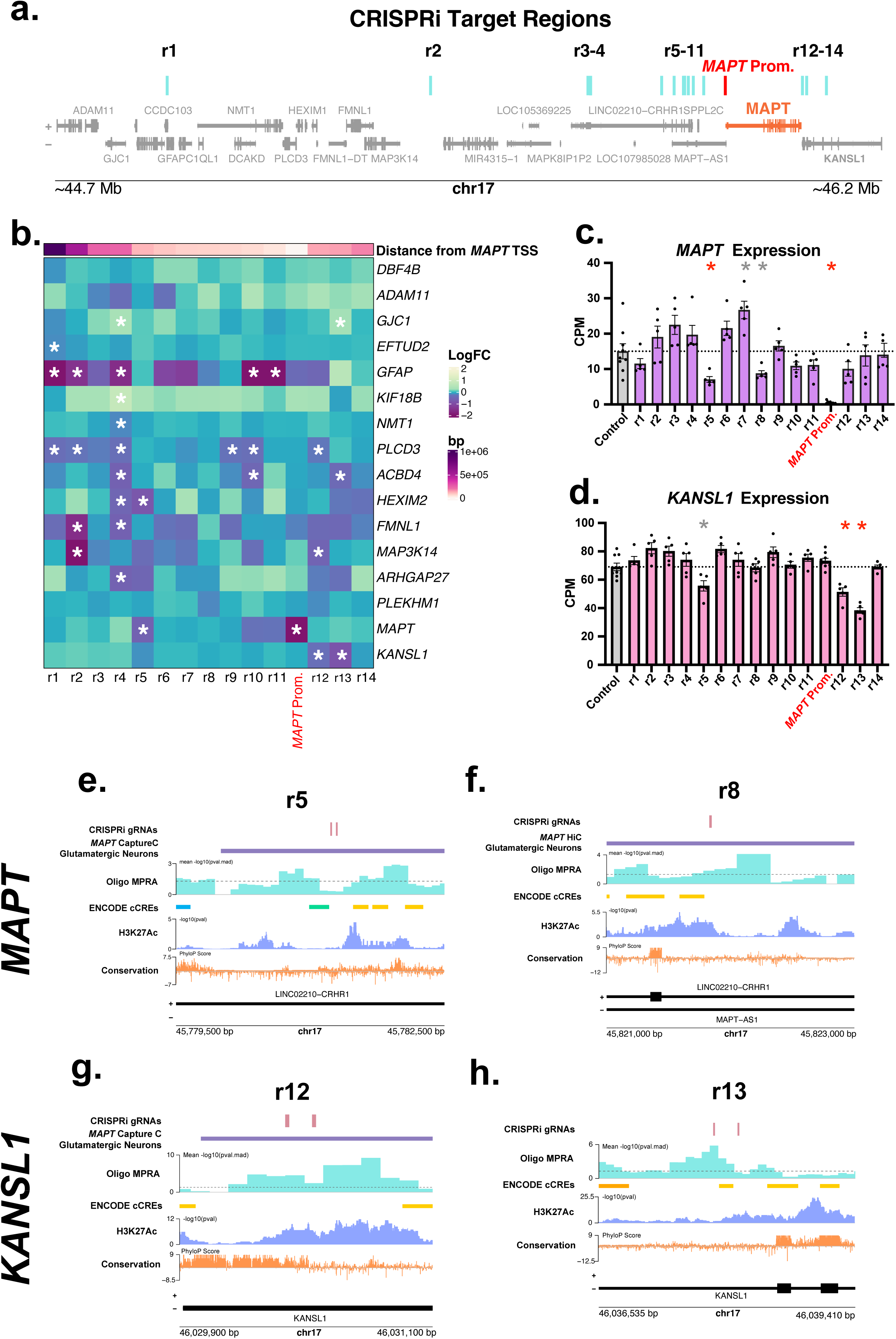
**a.** Gene track showing the locations of the nominated CRISPRi target regions in relation to the *MAPT* gene. Blue lines indicate target regions, and the red line is the location of the guide for the *MAPT* promoter positive control. **b.** Heatmap of genes 200kb away from a CRIPRi target region normalized for cell type score. Asterisks represent an adj. p. value ≤ 0.05. **c.** Barplot of *MAPT* CPM when targeting each nominated region with CRISPRi. Red asterisks represent targets with an adj. p. value ≤ 0.05 from DESeq2 both before and after normalizing for cell type score, gray asterisks represent an adj. P. value ≤ 0.05 from DESeq2 without normalizing for cell type score. **d.** Barplot of *KANSL1* CPM when targeting each nominated region with CRISPRi. Red asterisks represent targets with an adj. p. value ≤ 0.05 from DESeq2 both before and after normalizing for cell type score, gray asterisks represent targets with an adj. P. value ≤ 0.05 without normalizing for cell type score. **e–h.** Gene tracks of region r5, r8, r12, and r13 respectively. The dotted line on the MPRA signal track represents a pval.mad of 0.05.

### MPRAs identify regulatory variant effects at proximal CREs to MAPT

To determine the effects of genetic variation at CREs surrounding *MAPT*, the top 25% of CREs from the Oligo MPRA surrounding *MAPT* in neurons, as well as CREs previously identified for *MAPT*^39^ were selected as input into a variant screening MPRA. From the Alzheimer’s Disease Sequencing Project (ADSP) database, we selected single nucleotide variants (SNVs) and insertions/deletions (InDels) that overlapped these regulatory regions and were present in both Alzheimer’s Disease and non-Alzheimer’s Disease groups. We chose to exclude InDels greater than 10 bp. The MPRA was transfected into HEK293FT cells to generate lentivirus to transduce 14 day old KOLF2.1-hNGN2 neurons. DNA and RNA were collected from both cell types, and variant effects were quantified by bcalm^62^, with a |logFC| cut-off of 0.1 by comparing to reference sequence controls. The majority of variant effects were cell-type specific, and 2 InDel effects were shared between cell types (the deletions chr17:45853273:CAGGT:C (within *MAPT-AS1* intron 1), and chr17:46055010:CTGACTAA:C (within *KANSL1* intron 6)) (fig. 3a, tables s10-11). We identified 117 InDels showing a loss of activity in HEKs and 37 showing a gain of activity, as well as 5 showing a loss of activity and 9 showing a gain of activity in neurons (adj. p. value ≤ 0.05 and |logFC| > 0.1). SNVs showed a similar pattern with the majority of variant effects being cell-type specific (fig. 3b, tables s12-13). 55 SNVs showed an increase in activity and 61 a decrease in activity in neurons (adj. p. value ≤ 0.05 and |logFC| > 0.1). 447 and 433 SNVs showed a loss and gain of activity respectively, in HEKs, and 9 SNVs showed a loss of activity in both HEKs and neurons with 13 showing a gain of activity in both cell types. The SNVs with the largest effect sizes in neurons were chr17:46099360:C:A (within *KANSL1* intron 2) which increased activity of the CRE (adj. p. value = 0.0269), and chr17:46057315:T:C (within *KANSL1* intron 6) that showed a loss of activity (adj. p. value = 0.0246). Since the variants tested were chosen within regions that were CREs identified in neurons and not necessarily CREs in HEKs, that could contribute additional noise to the HEK variant effects. The candidate *MAPT* CRE region r8 showed a variant with a loss of activity at chr17:45821215:C:T in neurons (adj. p. value = 3.76×10^-5^) and gain of activity at chr17:45822291:C:G (adj. p. value = 0.0284) (fig. 3c). A CRE identified by Rogers et. al. at +77,758 bp from the *MAPT* promoter^39^ had 4 variant effects with chr17:45971571:TC:T and chr17:45971740:C:T showing a loss of activity and chr17:45971834:T:G and chr17:45971845:T:G showing a gain of activity in neurons (adj. p. values = 0.0221, 0.0495, 1.17×10^-5^, and 5.42×10^-3^ respectively) (fig. 3d). It should also be noted that this region also overlapped with regulatory regions in Cooper et al.^42^ and Corces et al.^63^

**Figure 3.**
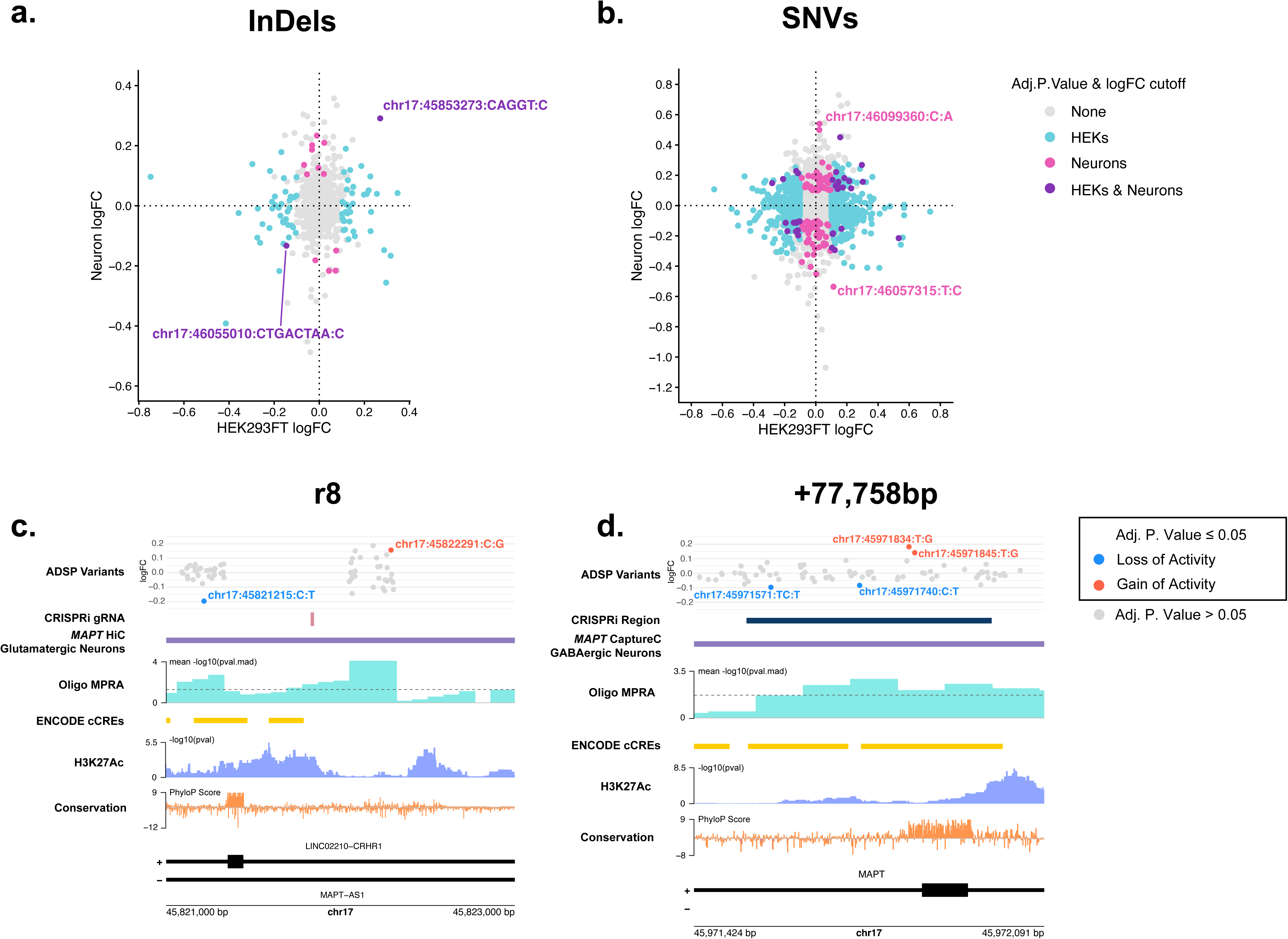
**a–b.** Correlation plots of (a) InDels (logFC Spearman’s ⍴ = 0.0649, approximate p = 0.114) and (b) SNVs (logFC Spearman’s ⍴ = 0.0865, approximate p = 1.52×10^-15^) variant effects from the ADSP database (both Alzheimer’s cases and unaffected controls). Blue, pink, and purple points represent variants with an adj. p. value ≤ 0.05 as well as a |logFC| > 0.1 in HEK293FT, Neurons, and both cell types respectively. Gray points represent variants that either have an adj. Pp value > 0.05 or a |logFC| < 0.1 **c–d**. Gene tracks showing (c) region r8 and (d) a CRE for *MAPT* identified by Rogers et al.^39^ respectively. The dotted line on the MPRA signal represents a pval.mad of 0.05. Variant effects are shown as logFC in neurons.

### Saturation mutagenesis comprehensively evaluates SNV regulatory effects at the MAPT promoter

To comprehensively assess the effect of genetic variation in the *MAPT* promoter, a 2,000 bp region encompassing the promoter was selected, and an MPRA was designed with 219 bp oligos centered on every base in this region (chr17:45,893,536–45,895,535). Oligos were designed with each possible alternate base relative to the reference sequence. To completely ablate any transcription factor binding sites, a centered 5 bp deletion sequence was also designed (fig. 4a). Activity signal from the initial Oligo MPRA in this region aligns with transcription factor binding sites in neurons (fig. 4b).^64^ This HOT (high occupancy target)^65,66^ site (chr17:45,893,810–45,894,854) was also enriched for single nucleotide regulatory variant effects with 332 of the 391 SNVs variant effects identified falling in this region (fig. 4b). Several variants in this region were present in the ADSP database, of note the common SNV rs11575896 (allele frequency = 0.1436, gnomAD v4.1.0^67^) shows an increase in activity that is specific to the G:A substitution (adj. p. value = 1.19×10^-3^) (fig. 4c). In addition, chr17:45894238:C:G showed an increase in enhancer activity in our study in neurons (logFC = 0.141, adj. p. value = 8.68×10^-3^) as well as in neural progenitor cells by Gaynor-Gillett et al.^43^ However, this variant had the opposite effect in HEKs (logFC = -0.0569, adj. p. value = 0.0414). Overall, we had 192 SNVs showing a loss of regulatory activity (adj. p. value ≤ 0.05 and logFC < -0.1), and 199 showing a gain of regulatory activity (adj. p. value ≤ 0.05 and logFC > 0.1). 142 and 82 centered 5 bp deletions had a loss and gain of activity, respectively (adj. p. value ≤ 0.05 and |logFC| > 0.1) (fig. 4b, table s14). In HEK293FT cells, we observed 523 SNVs and 417 SNVs with a gain and loss of activity, respectively, as well as 226 and 229 centered 5 bp deletions with a gain or loss of activity, respectively (adj. p. value ≤ 0.05 and |logFC| > 0.1) (fig. s3a, table s15).

**Figure 4.**
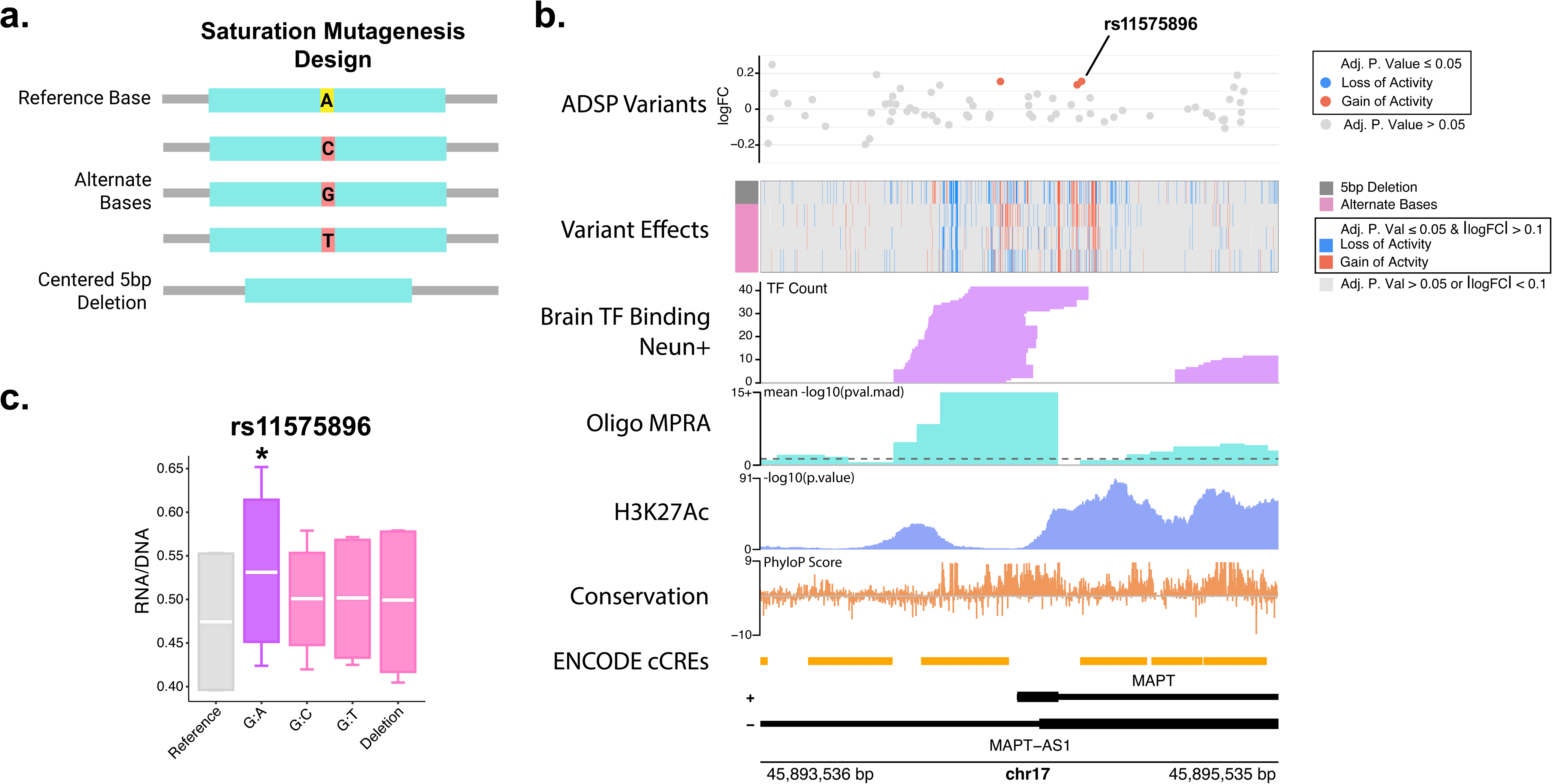
**a.** Graphical overview of *MAPT* promoter saturation mutagenesis design. **b.** Gene track for the *MAPT* promoter saturation mutagenesis region. The top track is variants found in ADSP in this region. Red points represent variants with an adj. p. val ≤ 0.05 with an increase of regulatory activity relative to the reference sequence. The second track is a heatmap showing variants that had an adj. p. val ≤ 0.05 as well as a logFC in activity of > 0.1 in red and < 0.1 in blue. Red lines indicate variants with an increase in regulatory activity, and blue lines indicate variants with a decrease in regulatory activity. The top row of the heatmap is data from the 5 bp deletions, the subsequent 3 rows are data from the 3 alternate bases at that position. The third track shows transcription factor binding in neurons, data from Loupe et al.^64^ The dotted line on the MPRA signal track indicates a pval.mad of 0.05. **c.** Box plot showing RNA/DNA ratio for rs11575896, the reference sequence, each alternate base tested, and 5 bp deletion at this region. Whiskers show maximum and minimum values. The asterisk indicates an adj. p. value ≤ 0.05 by bcalm.

### Correlation of AI predictive tools and MPRA results identify a high confidence loss of activity region in the MAPT promoter

Recently, there has been an increase in the use of AI tools in the interpretation of genomic datasets. We input the SNVs introduced in the saturation mutagenesis of the *MAPT* promoter into the AI tools AlphaGenome’s saturation mutagenesis function^68^and PromoterAI^69^, and compared the predicted regulatory effects on the *MAPT* gene with the activity of variant effects in our MPRA (tables s16-17). Overall, there was correlation between scores from AlphaGenome and PromoterAI (Spearmans’s ⍴ = 0.153, approximate p < 2.2×10^-16^) (fig. s4, fig. 5a). We compared the results of the saturation mutagenesis in both HEKs and neurons to determine variant effects specific to the neuron cell type, as well as variant effects shared between both cell types. Of the SNVs effects found in our *MAPT* promoter mutagenesis, 195 variant effects were shared between cell types (adj. p. value ≤ 0.05, |logFC| > 0.1 or < -0.1, with shared directionality of logFC in both cell types) and 186 were specific to neurons (fig. s3b, fig. 4b). Within the shared variant effects, there was a region at chr17:45,894,685–45,894,691 where any base substitution resulted in a gain of regulatory activity as measured by our MPRA (fig. 5b); however, this region shows a strong predicted loss of *MAPT* gene expression by both AlphaGenome and PromoterAI (fig. 5a). This region overlaps the *MAPT* exon1-intron1 boundary, and the variants are predicted to result in alternate splicing of the *MAPT* gene by SpliceAI^70^ (fig. 5c), accounting for the predicted loss of *MAPT* expression by the AI tools despite the increase in activity seen in the MPRA output. This underlies the importance of considering genomic context when interpreting results from MPRAs incorporating genetic variants. Among the neuron-specific variant effects, we observed there to be a region around chr17:45,894,271–45,894,297 where any base substitution resulted in a loss of regulatory activity (fig. 5d). This region overlaps an area of strong activity signal from our initial MPRA as well as an area of high transcription factor binding (fig. 4b). The peaks of loss of activity correspond to peaks of high genetic conservation scores^71,72^, and we believe this to be a critical transcription factor binding site for *MAPT* expression in neurons. MotifbreakR^73^ was used to determine what transcription factor binding sites were disrupted by the variants at these positions. We saw that in this region binding sites were disrupted for the transcription factors TCFL5, EGR2, and ZBTB14 (fig. 5e–f, table s20). When comparing the scores given by AlphaGenome and PromoterAI with the results from the MPRA, we saw that the neuron-specific loss of activity region (in fig. 5d) was the region of highest agreement among all three tools, while the exon1-intron1 boundary region (in fig. 5b) was the area of highest disagreement among the MPRA results and AI tools (fig. 5g).

**Figure 5.**
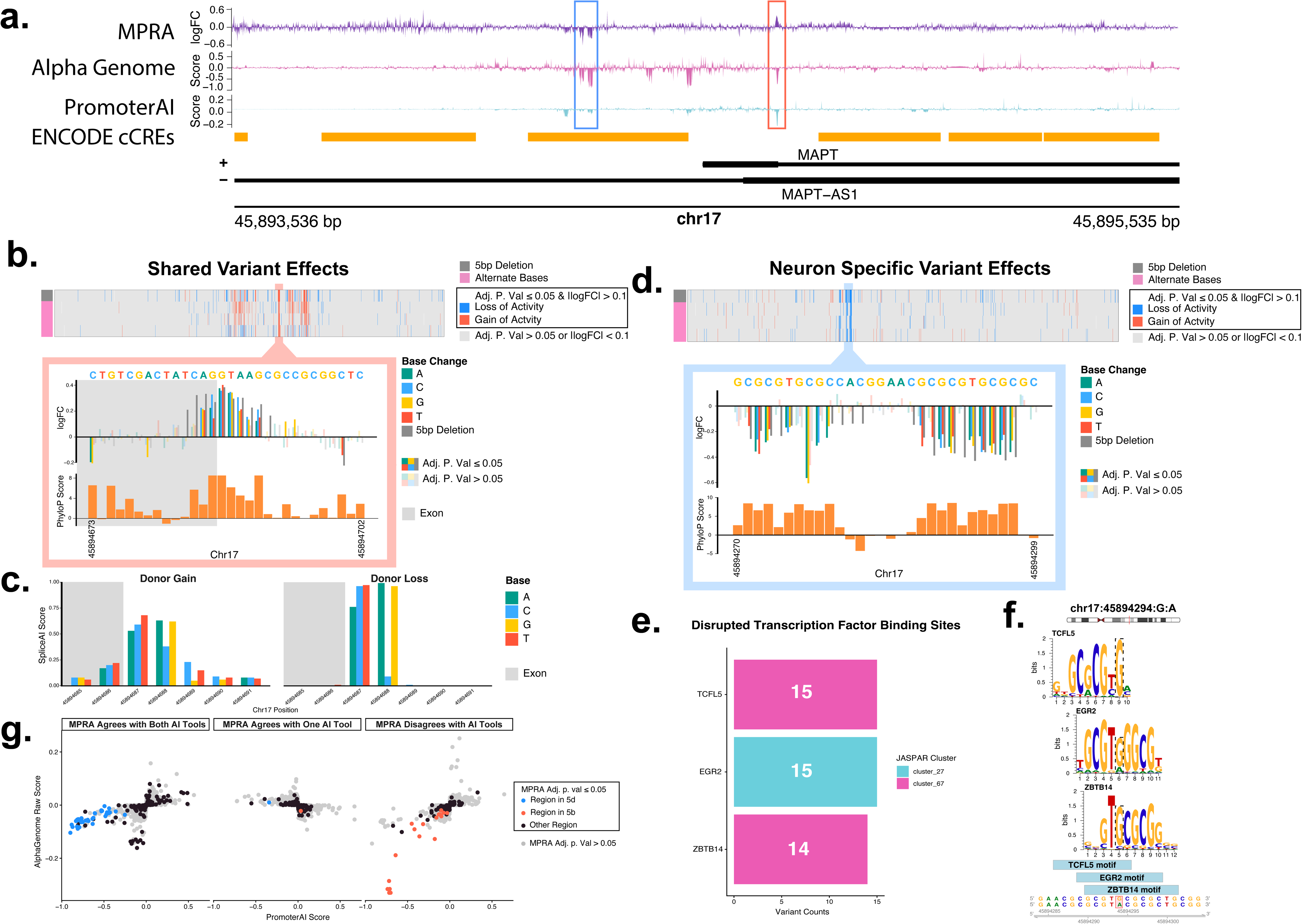
**a.** Gene track showing logFC of each SNV tested in the *MAPT* promoter region (top), AlphaGenome score for each variant in this region in glutamatergic neurons (middle), and PromoterAI (bottom) score for each variant in this region. The blue box represents the region in 5d and the red box represents the region in 5b **b.** (top) Heatmap showing variants with an adj. P. value ≤ 0.05 by bcalm and a |logFC| > 0.1 that are shared between neurons and HEK293FT cells. The top row of the heatmap shows data from 5 bp centered deletions at that position, the bottom 3 rows show data from alternate bases at that position. Variants showing an increase in regulatory activity compared to reference are shown in red and variants with a decrease in regulatory activity are shown in blue. (bottom) logFC of regulatory activity of each variant in the highlighted region compared to the reference sequence. Opaque bars have an adj. P. value ≤ 0.05 by bcalm and transparent bars have an adj. P. value > 0.05. Bars with a gray background are in exon 1 of *MAPT* and bars with a white background are in intron 1 of *MAPT*. Reference sequence is shown on top of the graph. Below the top graph is a bar plot of PhyloP genetic conservation score^71,72^ for each base. **c.** Bar plots showing SpliceAI scores for each alternate base in this region. Bars with a gray background are in exon 1 of *MAPT* and bars with a white background are in intron 1 of *MAPT*. **d.** (top) Heatmap showing variants with an adj. P. value ≤ 0.05 by bcalm and a |logFC| > 0.1 that are present in neurons but not in HEK293FT cells. The top row of the heatmap shows data from 5 bp centered deletions at that position, the bottom 3 rows show data from alternate bases at that position. Variants showing an increase in regulatory activity compared to reference are shown in red and variants with a decrease in regulatory activity are shown in blue. (bottom) logFC of regulatory activity of each variant in the highlighted region compared to the reference sequence. Opaque bars have an adj. P. value ≤ 0.05 by bcalm and transparent bars have an adj. P. value > 0.05. Bars with a gray background are in exon 1 of *MAPT* and bars with a white background are in intron 1 of *MAPT*. Reference sequence is shown on top of the graph. Below the top graph is a bar plot of PhyloP genetic conservation score^71,72^ for each base. **e.** Transcription factor binding sites that are disrupted by variants with a significant loss of activity in the highlighted region in d. in motifbreakR colored by JASPAR motif clusters. **f.** Precision weight matrices of disrupted transcription factor binding sites in the highlighted region in d. Pictured is the base chr:45894294:G:A which is one of the variants with a significant loss of activity in the highlighted region in d. **g.** Correlation plots of AlphaGenome and PromoterAI scores separated by whether the direction of the predicted variant effect agrees with the direction of the logFC seen in the MPRA. Gray dots indicate an adj. p. value > 0.05 by bcalm. Black points indicated an adj. P. value ≤ 0.05 not within a highlighted region. Blue points indicate regions with an adj. P. value ≤ 0.05 in the highlighted region in d. Red points indicate regions with an adj. P. value ≤ 0.05 in the highlighted region in b.

## Discussion

We screened a ∼3 Mb region around the *MAPT* locus for cis-regulatory elements in human iPSC derived excitatory neurons using lentiMPRAs. The results of these screens identified several previously unannotated regulatory elements and also validated activity of existing ENCODE cCREs. Our results show the importance of testing CREs in cell types of interest, as the majority of regulatory elements we identified were cell-type specific. We believe this also accounts for the findings of multiple new previously unannotated CREs in this region. Using known genetic variation from Alzheimer’s disease patients and non-Alzheimer’s disease affected individuals from the ADSP database, we were able to identify variants that altered the activity of the CREs with the highest activity around *MAPT.* These variants likewise showed strong cell type specificity. We identified regulatory variant effects at CREs for *MAPT* identified in Rogers et al. at regions -674,458; -652,338; -464,677; -461,949; -44,905; -20,178; and +77,758 bp away from the *MAPT* promoter.^39^ Due to their location within validated CREs for *MAPT*, these variants have the potential to alter *MAPT* expression although their direct effect on *MAPT* was not tested in this study.

We comprehensively mapped genetic variant effects at a 2,000 bp region encompassing the *MAPT* promoter (chr17:45,893,536–45,895,535) and identified 199 variants with an increase in regulatory activity and 192 variants with a decrease of regulatory activity relative to the reference genome in neurons. Of note, we identified 145 neuron-specific single nucleotide regulatory variants that were located in a HOT site region of the *MAPT* promoter (chr17:45,893,810–45,894,854) shown by strong MPRA activity and transcription factor binding in neurons that have an increased likelihood to alter expression of *MAPT*, including a variant also found by Gaynor-Gillett et al. in neural progenitor cells.^43^ We also identified a ∼26 bp region at chr17:45,894,271–45,894,297 with a high confidence reduction in regulatory activity (fig. 5d). Using motifbreakR, we saw that this region was enriched for the binding motifs of the transcription factors EGR2, TCLF5, and ZBTB14. EGR2 is an immediate early gene (IEG) heavily implicated in promoting myelination^74–78^ and Charcot-Marie Tooth Disease^79–89^ as well as being implicated in memory, epilepsy, inflammatory response, and Rett syndrome.^82–89^ Interestingly, *EGR2* was recently shown to be upregulated in the hippocampus of a presymptomatic rat model of AD, and *EGR2* deficient mice show an improvement in learning and memory.^82,90^ ZBTB14 is implicated in seizure regulation^91^, and TCLF5 has been studied in spermatogenesis.^92,93^ It is worth noting that both EGR2 and ZBTB14 were targeted in a genome wide CRISPRi screen in human iPSC-derived neurons by Samelson et. al. where no effect was observed on levels of Tau oligomers.^94^ Likewise, EGR2 was tested in shRNA screens in human Daoy cells as well as in drosophila and with no change in Tau protein levels.^95^ Since neither of these papers measured gene expression, it is possible that these transcription factors modulate *MAPT* gene expression in a way that is not translated to protein expression levels. The transcription factors could also be acting in a cell-type and species specific way, and since we did not directly observe the effect of these transcription factors on *MAPT* expression it is possible that other transcription factors with similar binding motifs are responsible for the loss of activity at this region observed in our MPRA. In addition, this is not the only region of the *MAPT* promoter responsible for inducing *MAPT* transcription. The *MAPT* promoter itself is a genomic HOT site with multiple transcription factors binding this region (fig. 4b), including CLOCK, EGR1, and NRF1 (table s18).^64^

With the advent of new AI tools to predict the regulatory effects of genetic variation, we compared the results from our saturation mutagenesis MPRA with two AI tools, PromoterAI and AlphaGenome. Overall, there was a correlation between two tools (Spearmans’s ⍴ = 0.153, approximate p < 2.2×10^-16^). However, PromoterAI had a higher correlation with the results from our MPRA (Spearmans’s ⍴ = 0.482, approximate p = 1.89×10^-4^ PromoterAI, and Spearmans’s ⍴ = 0.0277, approximate p = 0.0321 AlphaGenome). We observed that the region with the highest agreement among all three tools was also the region of largest variant effects measured by our MPRA. It is worth noting that the MPRAs do not reflect direct regulation of the *MAPT* gene, only accounting for isolated increases or decreases in regulatory activity of the tested sequences in each cell type. The AI tools strive to predict expression effects of specific genes by pulling in multiple predictive datasets, accounting for some of the differences between our MPRA and the AI tools. We believe that the field is not yet at a point where MPRAs can be excluded altogether in favor of AI predictive tools due to the highly cell type specific nature of gene regulation, but they can aid in the interpretation of MPRA datasets as we saw with the splice variant region that was in high agreement between both AI tools but showed an increase in activity in our saturation mutagenesis MPRA.

Limitations of this study include that all experiments were done in cell types containing the H1 haplotype, and it is possible that experimental results would be different in H2 containing cells. In addition, the direct effect of genetic variation on *MAPT* expression was not evaluated at this time, and is beyond the scope of this study (requiring techniques such as prime editing, which could be the focus of future studies). Additionally, the only cell types used in this study were excitatory neurons from the KOLF2.1Jh-NGN2 line, HEK293FT cells, or mixed KOLF2.1J neuron cultures and other cell types were not tested. Despite these limitations, this study makes strong advances in understanding the effects of regulatory elements and related genetic variation on expression of *MAPT* and other genes near *MAPT* also affected by the H1/H2 inversion event associated with several neurodegenerative diseases.

## Methods

### BAC MPRA

#### Library Design

24 BACs clones (fig. s1, table s1) were selected from BacPac Resources (https://bacpacresources.org) that aligned to a region approximately 3 Mb around the *MAPT* locus. BACs were grown in 5mL Terrific Broth (Corning 46-055-CM) with 12.5µg/mL (sigma C0378-5G). 5µg of BAC DNA was purified using QIAGEN plasmid buffer set (QIAGEN 19046) and QIAGEN-tip 500 (QIAGEN 10063), and sheared to an average size of around 250 bp using a Covaris S220. Sheared BACs were prepped for cloning as previously described^96^, and they were inserted into pLS-SceI following the published protocol.^97^ Following cloning and electroporation into NEB 10-beta electrocompetent cells (New England Biolabs C3020K), a 500mL LB broth (Corning 46-050-CM) + 100µg/mL carbenicillin (Sigma C1389) culture was grown at 37°C and 150rpm for 13 hours. The plasmid pool was purified using the QIAGEN plasmid plus mega kit (QIAGEN 12981). A subset of the culture was spread on LB agar plates (Millipore 1102830500) with 100µg/mL carbenicillin (Sigma C1389) at 37°C, and several colonies were chosen for sequence validation with Sanger Sequencing before sending the pool for Barcode Association Sequencing. pLS-SceI was a gift from Nadav Ahituv (Addgene plasmid # 137725; http://n2t.net/addgene:137725; RRID:Addgene_137725)^97^.

#### Barcode Association

The locus was divided into 100 bp nonoverlapping bins. The sheared BAC sequences were aligned to HG38, and a custom script was used to aid in the association of each 15 bp barcode with a 100 bp bin based off of which bin the middle of the associated sheared BAC sequence was aligned. Libraries were prepared according to Gordon et al^97^ and sequenced on an Illumina NextSeq flowcell (Table s21).

### Oligo MPRA

#### Library Design

270 bp oligos were ordered from Twist biosciences every 90 bp for the forward and reverse DNA sequences of regions of interest (ROI). Total, this gives 6X coverage of each ROI tested in each cell type. For analysis, forward and reverse strands were combined into 1 ROI. Regions were chosen based on areas with low coverage in the BAC MPRA, tiling the *MAPT* locus (Hg38 chr17:45,771,561–46,100,061), as well as further upstream (Hg38 chr17:44,400,964–44,575,474 and Hg38 chr17:44,605,938–44,718,798). Several regions were included for other genes of interest based on multiomics data from previous publications (Anderson and Rogers et al.^52^ and Cochran et al.^98^), and 100 scrambled sequences were added as negative controls.

The oligo pool was inserted into pLS-SceI according to the previously published protocol.^97^ Following cloning and electroporation into NEB 10-beta electrocompetent cells (New England Biolabs C3020K), a 500mL LB broth (Corning 46-050-CM) + 100µg/mL carbenicillin (Sigma C1389) culture was grown at 37°C and 150rpm for 13 hours. The plasmid pool was purified using the QIAGEN plasmid plus mega kit (QIAGEN 12981). A subset of the culture was spread on LB agar plates (Millipore 1102830500) with 100µg/mL carbenicillin (Sigma C1389) at 37°C, and several colonies were chosen for sequence validation with Sanger Sequencing before sending the pool for Barcode Association Sequencing on an Illumina Novaseq (Table s21).

### Variation MPRA

#### MAPT Promoter Mutagenesis

219 bp oligos were designed for a 2,000 bp region surrounding and including the *MAPT* promoter (Hg38 chr17:45,893,536–45,895,535) and ordered from Twist Biosciences. For every base in this region, an oligo was designed where the reference sequence is centered on the position of interest as well as 3 additional oligos with each possible alternate base at that position and an oligo with a centered 5 bp deletion. The mutagenesis and known variant MPRAs were combined in a single 36,000 Oligo MPRA pool and separated at the time of analysis.

#### Known Variants

SNVs and InDels of no more than 10 bp were chosen from the ADSP^13^ database that overlapped the top 25% of CREs in neurons near *MAPT* (Hg38 chr17:45,771,561–46,100,061) found using the Oligo MPRA, regions chosen for CRISPRi testing from the BAC or Oligo MPRA, the *MAPT* promoter (Hg38 chr17:45,893,536–45,895,535), and regions that were shown as CREs for *MAPT* (+100 bp on each side) from a previous publication (Rogers et al.^39^). For SNVs, 219 bp oligos were designed with the variant centered for both the reference sequence and the SNV. For deletions, the reference seq was 219 bp for deletions of an odd number of bases and 218 bp for deletions of an even number of bases in order to center the deletion region. The variant sequences were the length of 219 or 218 bp minus the number of bases being deleted. For insertions, the variant sequence was 219 or 218 bp (depending on if it was an even or odd number of bases inserted) in order to have the inserted sequence centered within the reference sequence. Reference sequences were 219 or 218 bp long minus the number of bases inserted in the alternate allele. Variants with an allele count of 0 were excluded and variants with an allele frequency of 1 or more were represented 2X. A total of 627 InDels and 9,090 SNVs were included. Since the majority of included regions were previously positive in an MPRA, reference sequences for these regions can act as positive controls.

12 promoters were included as additional positive controls and were represented 2X. The scrambled controls were included from the Oligo MPRA, as well as 2,510 MPRA negative regions and 2,539 GC-matched scrambled negative controls from the tested regions. A subset of negative controls were shortened so that there was a length match to each length of tested regions in the oligo pool, as well as altering the middle base of the scrambled region to each alternate base. The oligo pool was ordered from Twist biosciences. Plasmid libraries were prepared as in Gordon et al.^97^ with a slight modification as the protocol was modified as previously described in order to test both orientations of the insert.^99^ Following cloning into pLS-SceI and electroporation into NEB 10-beta electrocompetent cells (New England Biolabs C3020K), a 500mL LB broth (Corning 46-050-CM) + 100µg/mL carbenicillin (Sigma C1389) culture was grown at 30°C and 150rpm for 20 hours. The plasmid pool was purified using the QIAGEN plasmid plus mega kit (QIAGEN 12981). A subset of the culture was spread on LB agar plates (Millipore 1102830500) with 100µg/mL carbenicillin (Sigma C1389) at 37°C overnight, and several colonies were chosen for sequence validation with Sanger Sequencing before sending the pool for Barcode Association Sequencing on an Illumina NovaSeq X Plus (Table s21).

### MPRAs

We followed the LentiMPRA protocol.^97^ Briefly, Kolf 2.1J-NGN2 iPSCs were obtained from Jackson laboratory (JIPSC002070). iPSCs were grown as aggregates in matrigel coated flasks in MTESR Plus media (Corning 354277, StemCell Technologies 100-0276). Cells were treated with accutase (Stemcell Technologies 07920) and were plated as single cells onto 15cm plates coated with both poly-l-ornithine (Sigma P4957) and Matrigel (Corning 354277) at a density of 25,000 cells/cm^2^ in induction media (DMEM/F12 with HEPES Gibco 11039021, 1X N2 Supplement-A Stemcell Technologies 07152, 1X Glutamax ThermoFisher 35050061, 1X MEM non-essential amino-acids Gibco 11140050, and at plating 10 μM ROCK inhibitor (Y-27632) (Stemcell technologies 72304)). Doxycycline (Stemcell Technologies 72742) was added to the media at a concentration of 2µg/mL to induce *NGN2* expression and differentiation into excitatory neurons. The cell media was swapped to cortical neuron media (BrainPhys Neuronal Medium Stemcell Technologies 05790, 1X NeuroCult SM1 Neuronal Supplement Stemcell Technologies 05711, 5µg human recombinant BDNF Stemcell Technologies 78005, 5µg human/mouse recombinant NT-3 Stemcell Technologies 78074, 0.5mg Laminin Mouse Protein Gibco 2307015) and allowed to differentiate for 14 days. 1 hr before transduction with lentivirus, the media was swapped to cortical neuron media containing protamine sulfate (Sigma-Aldrich P3369) at a concentration of 10 µg/mL. Lentivirus was made by transfecting HEK293FT cells plated at a density of 73,000 cells/cm^2^ in a T225 flask in supplemented DMEM media (Gibco 11965-084, 10% FBS Cytivia SH30071.03, 1X Glutamax ThermoFisher 35050061, 1X MEM non-essential amino-acids Gibco 11140050, Geneticin 50µg/mL Gibco 10131035). The following day, the media was swapped to Optimem + Glucose (Gibco 31985070, 0.6mg/L Sigma G8644, 1X Glutamax ThermoFisher 35050061). 2 days later the media from 2 T225 flasks per biological MPRA replicate was filtered at 0.45µm (ThermoFisher 09-740-63A). 11.25mL of virus was added to each 15cm plate of neurons. 24hrs following transduction, the media was swapped to fresh cortical neuron media. Cells were collected on day 18 using the Qiagen DNA/RNA kit and on-column DNAse digestion (QIAGEN 80204 and 79254). DNA and RNA from 2 15cm plates were combined for each biological replicate of the MPRAs. Libraries were prepared as in Gordon et al.^97^ and sequenced on a mixture of Illumina NovaSeq X Plus (Variant MPRA, Oligo MPRA, BAC MPRA), Illumina Novaseq (Oligo MPRA), and Illumina NextSeq (BAC MPRA) flowcells (Table s21). All mammalian cells were grown at 37°C and 5% CO_2_.

#### MPRA Analysis

For the Oligo MPRA, the barcode association dictionary was generated with MPRAflow.^97^ The BAC MPRA and Oligo MPRA both used MPRAflow^97^ to generate DNA and RNA barcode counts. MPRAnalyze^100^ was used to analyze both MPRAs. For the variant and mutagenesis MPRAs, barcode association dictionaries were generated with MPRAflow^97^ without a mapping quality filter. Oligos of different lengths were associated with barcodes separately using a CIGAR score filter requiring a perfect match according to the length (219M, 218M, etc.). The barcode dictionaries were combined, and barcodes associating with multiple oligos were removed. Counts of each barcode in the RNA and DNA fractions from each cell type were generated with MPRAsnakeflow^101^ and input into bcalm.^62^ Known InDels, SNVs, and the promoter mutagenesis were analyzed separately. InDels and SNVs alternate sequences were compared to reference sequence controls. InDels and SNVs were analyzed separately to avoid any effect of altering oligo sizes in the InDels to affect the model for SNV analysis. For the saturation mutagenesis, each alternate base as well as the 5 bp deletion were compared to the reference sequence for that region. Both orientations of test sequences were combined into 1 region for analysis. motifbreakR^73^ was used to identify transcription factor binding sites. SpliceAI^70^ scores were accessed from https://spliceailookup.broadinstitute.org.

### CRISPRi

gRNAs were designed for each region of interest using Benchling and were ordered from IDT (table s6). The guides were cloned into the pLV hU6-sgRNA hUbC-dCas9-KRAB-T2a-Puro vector^102^ (addgene 71236) using FASTAP (ThermoFisher EF0651) and ESP3I and in 10X fast digest buffer (ThermoScientific FD0454). Lentivirus was created by transfecting HEK293FT (Invitrogen R70007) cells plated at 700,000 cells/well in a PLO (Sigma 00107) coated 6 well plate in supplemented DMEM media (as above) and cells were transfected with 0.75µg psPAX2 (Addgene 12260), 0.5µg pMD2.G (Addgene 12259), and 0.5µg pLV hU6-sgRNA hUbC-dCas9-KRAB-T2a-Puro (Addgene 71236) containing each gRNAs to each target region of interest with Lipofectamine LTX (Invitrogen 15338100). Kolf2.1 NPCs were generated as previously described.^39^ and plated at a density of 200,000 cells per well in a matrigel coated (Corning 354230) 12 well plate in NPC media (350mL DMEM Gibco 11995081, 150mL Ham’s F12 Gibco 11765054, 1X B-27 Supplement Gibco 17504044, 40ng/mL bFGF R&D Systems 233-FB, 20ng/mL hEGF Sigma E9644, heparin 5µg/mL Sigma H3149). The following day, media from the HEK293FT cells was filtered using a 0.45µm PES syringe filter (VWR 76479-020) to obtain CRISPRi packaged lentivirus. Protamine sulfate 10µg/mL (Sigma-Aldrich P3369) was added to the NPC media at a concentration of 10 µg/mL 1 hour before transduction, and NPCs were transduced with 500µL of virus per well (250µL of guide 1 and 250µL of guide 2 lentivirus were added to each replicate, for region r8 500µL of guide 1 was added). 2 guides per well were used with the exception of region r8 where only one guide was tested due to sequence content of the target region and difficulties designing and cloning site specific guides for this region. pLV hU6-sgRNA hUbC-dCas9-KRAB-T2a-GFP (addgene 71237) was used in a separate well as a control. The next day, media was swapped on the NPCs to media + puromycin (0.5µg/mL) (Gibco A1113803) to select transduced cells. 48hrs after selection, the media on the NPCs to Brain Phys Differentiation media (BrainPhys Neuronal Medium Stemcell Technologies 05790, 1X N2 Supplement-A Stemcell Technologies 07152, 1X NeuroCult SM1 Neuronal Supplement Stemcell Technologies 05711, 10µg human recombinant GDNF Stemcell Technologies, 10µg human recombinant BDNF Stemcell Technologies 78005, 500µL of 200µM L-Ascorbic acid Sigma A0278, 0.5mg Laminin Mouse Protein Gibco 2307015) following the previously published Bardy et al. protocol.^39,53^ After 14 days of differentiation, the cells were collected using the Norgen RNA purification kit (Norgen 17200 and 25710). RNA was sent to Novogene for mRNAseq at 20 million paired end reads per sample on an Illumina NovaSeq X Plus. All mammalian cells were grown at 37°C and 5% CO_2_. pLV hU6-sgRNA hUbC-dCas9-KRAB-T2a-Puro was a gift from Charles Gersbach (Addgene plasmid # 71236; http://n2t.net/addgene:71236; RRID:Addgene_71236).^102^ pLV hU6-sgRNA hUbC-dCas9-KRAB-T2a-GFP was a gift from Charles Gersbach (Addgene plasmid # 71237; http://n2t.net/addgene:71237; RRID:Addgene_71237).^102^ psPAX2 was a gift from Didier Trono (Addgene plasmid # 12260; http://n2t.net/addgene:12260; RRID:Addgene_12260). pMD2.G was a gift from Didier Trono (Addgene plasmid # 12259; http://n2t.net/addgene:12259; RRID:Addgene_12259).

#### RNAseq

Cutadapt^103^ was used to trim adapters from reads before aligning them to GENCODE v42 using STAR aligner.^104^ The reads were indexed using Samtools^105^ and deduplicated using picard^106^. Reads were counted using htseq-count^107^. CPMs were generated using edgeR.^108^ To account for the varying numbers of cell types in the culture, a cell type score was assigned to each replicate using Seurat^54^ and a list of gene markers: neurons (*RBFOX3*, *SYP*), NPCs (*NES*, *PAX6*), and astrocytes (*ALDH1L1*, *SLC1A2*, *SLC1A3*, *S100B*). DESeq2^55^ was used to analyze each target region against a guide targeting the AAV Safe Harbor Locus^109,110^ filtering for a CPM of at least 1 with and without including cell type scores in the model.

### AI tools

#### AlphaGenome MAPT Suturation Mutagenesis

The AlphaGenome API was accessed via the DnaClient using Google Colab.^111^ Hg38v46 annotations were used as reference and a 1 Mb context window around *MAPT* was constructed for predicting variant effects. A saturation mutagenesis window encompassing a window 1kb upstream and downstream of the *MAPT* TSS was built. In silico mutagenesis was then conducted using the recommended RNAseq variant scorer and AlphaGenome’s built in saturation mutagenesis function. The effect of each SNV on RNAseq signal for each gene and pseudogene within the context window was compared against the effect of the reference allele as described by Avsec et al.^68^ resulting in raw scores. Raw scores predicting gene expression changes for *MAPT* in glutamatergic neurons (CL:0000679) were subsequently extracted and exported for analysis in RStudio.

#### PromoterAI MAPT Mutagenesis

A file representing all possible SNVs within 1kb upstream and downstream of the *MAPT* TSS was generated as specified by Jaganathan et al.^69^ Using an input sequence length of 20,480 bp centered around each variant, PromoterAI scored the effect of each SNVs on gene expression using the promoterAI_v1_hg38_mm10_finetune model and GCF_000001405.26 RefSeq. Variant scores were appended to each SNV and imported into RStudio for analysis.

## Graphing

Plots were made using ggplot2^112^, plotgardener^113^, complexheatmap^114^, or Graphpad prism 10. Graphical figures were made in BioRender and Adobe Illustrator.

## External Data

Known genetic variants were acquired from the Alzheimer’s Disease Sequencing Project (ADSP) (NIAGADS NG00067). Transcription factor binding and H3K27ac ChIP-seq data was generated by Loupe et al.^64^ and is available through the PsychENCODE Consortium at https://doi.org/10.7303/syn51942384.1. Brain single nucleus multiomics data was generated by Anderson et al.^52^ and is available at GEO accession number GSE214637. *MAPT* glutamatergic neuron HiC, *MAPT* glutamatergic neuron CaptureC, *MAPT* gabaergic cell Capture C, and cultured neuron single nucleus multiomics data were generated by Rogers et al.^39^ and are available at GSE228121. PhyloP conservation scores were generated by the Zoonomia Consortium^71,72^ and obtained from https://hgdownload.cse.ucsc.edu/goldenpath/hg38/cactus241way/. ENCODE cCREs were generated by Moore et al.^51^ and are available on SCREEN (https://screen.wenglab.org).

## Abbreviations

dELS: distal enhancer like signature
pELS: proximal enhancer like signature
TF: transcription factor signal
CA-CTCF: chromatin accessible, CTCF
CA: chromatin accessible
PLS: promoter like signature
CA-TF: chromatin accessible, transcription factor signature
CA-H3K4me3: chromatin accessible, H3K4me3 signal
MPRA: Massively Parallel Reporter Assay
CRE: Cis-regulatory Element
logFC: log(fold change)
HEKs: HEK293FT cells
AI: artificial intelligence

## Acknowledgements

This work was supported by NIH grants K99AG068271/R00AG068271 and R01AG085357 awarded to J.N.C. as well as by donors to the HudsonAlpha Foundation Memory and Mobility Program. This study used data generated by NIAGADS and the Alzheimer’s Disease Sequencing project (complete acknowledgments statement in extended acknowledgments section). We thank Jane Grimwood and the HudsonAlpha Genome Sequencing Center for their assistance in sequencing MPRA libraries.

## Author Contributions

Conceptualization, J.N.C; methodology, B.A.M.; Experimental Design, J.N.C and R.M.H.; Investigation, R.M.H., J.N.B., S.N.L., E.A.B., S.Q.J., B.B.R., J.W.T., and H.L.L.; writing—original draft, R.M.H. and J.N.C.; writing—review & editing, B.A.M., J.N.B., S.N.L., S.Q.J., B.B.R., H.L.L., J.W.T. and J.N.C.; funding acquisition, J.N.C.; resources, J.N.C.; supervision, J.N.C.

## Resource Availability

Further information and requests for resources and reagents should be directed to and will be fulfilled by the lead contact, J. Nicholas Cochran.

## Data and Code Availability

Processed and raw data generated in this study are available through NCBI GEO accession GSE325670. All original code generated for this study is available at https://github.com/HudsonAlpha/Hauser_MAPT_MPRAs.

## Figure Legends

**Figure s1.**
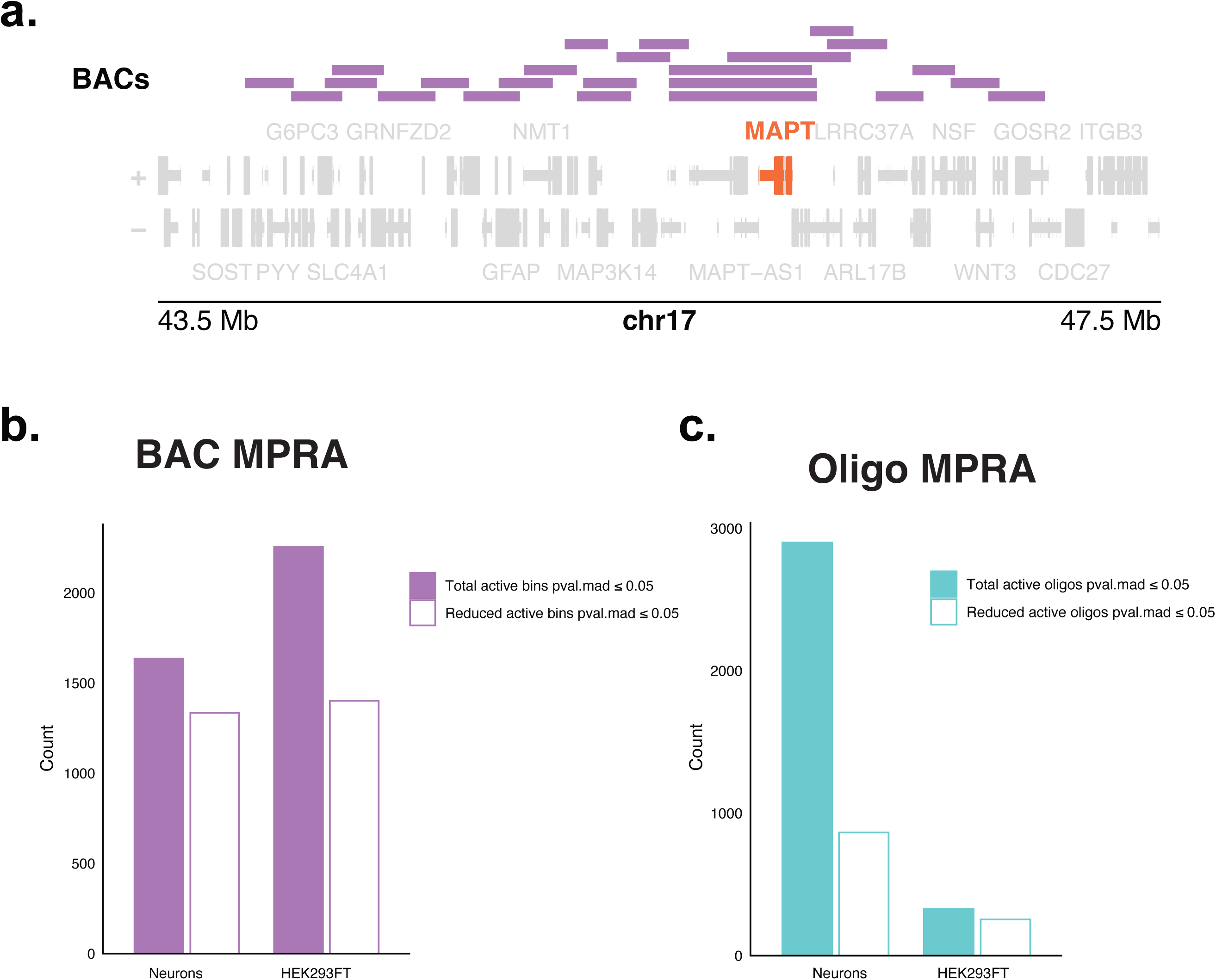
**a.** Gene track showing alignment of the 24 BACs chosen for the BAC MPRA to hg38. **b**. Bar plot showing the total number of bins with a pval.mad ≤ 0.05 in the BAC MPRA for Neurons and HEKs, as well as a reduced count where consecutive bins were merged into a single active region. **c.** Bar plot showing the total number of oligos with a pval.mad ≤ 0.05 in the Oligo MPRA, as well as a reduced count where consecutive and overlapping oligos were merged into a single active region.

**Figure s2.**
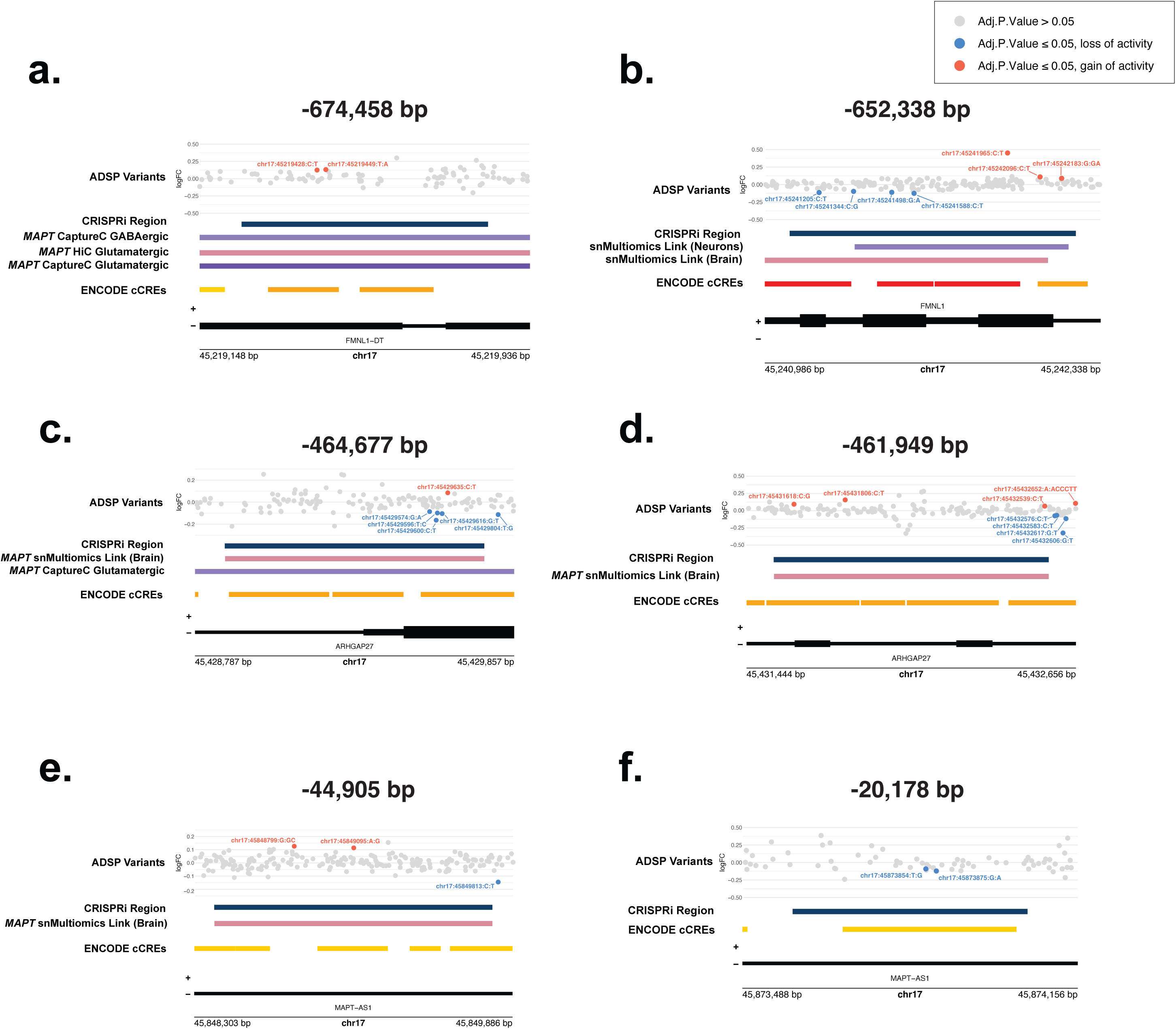
**a–f.** Gene tracks of CREs for *MAPT* identified in Rogers et al.^39^ where variant effects were found in our MPRA. Regions are labeled as in Rogers et al.^39^ as distance from the *MAPT* promoter. Variant effects shown are in Neurons. Adj. p. values for variants are: **a.** chr17:45219428:C:T = 0.0456, chr17:45219449:T:A = 0.0401, **b.** chr17:45241205:C:T = 0.0467, chr17:45241344:C:G = 0.0246, chr17:45241498:G:A = 8.16×10^-4^, chr17:45241588:C:T = 3.74×10^-4^, chr17:45241965:C:T = < 2.2×10^-16^, chr17:45242096:C:T = 0.0246, chr17:45242183:G:GA = 0.0165, **c.** chr17:45429574:G:A = 0.0175, chr17:45429600:C:T = 5.82×10^-4^, chr17:45429596:T:C = 3.17×10^-13^, chr17:45429616:G:T = 2.82×10^-5^, chr17:45429635:C:T = 2.04×10^-3^, chr17:45429804:T:G = 2.64×10^-3^, **d.** chr17:45431618:C:G = 0.0222, chr17:45431806:C:T = 1.17×10^-4^, chr17:45432539:C:T = 0.0476, chr17:45432576:C:T = 7.80×10^-3^, chr17:45432583:C:T = 0.0255, chr17:45432606:G:T = < 2.2×10^-16^, chr17:45432617:G:T = 1.88×10^-4^, chr17:45432652:A:ACCCTT = 4.32×10^-3^, **e.** chr17:45848799:G:GC = 0.0312, chr17:45849095:A:G = 1.88×10^-4^, chr17:45849813:C:T = 5.08×10^-3^, **f.** chr17:45873854:T:G = 7.38×10^-3^, chr17:45873875:G:A = 0.0237.

**Figure s3.**
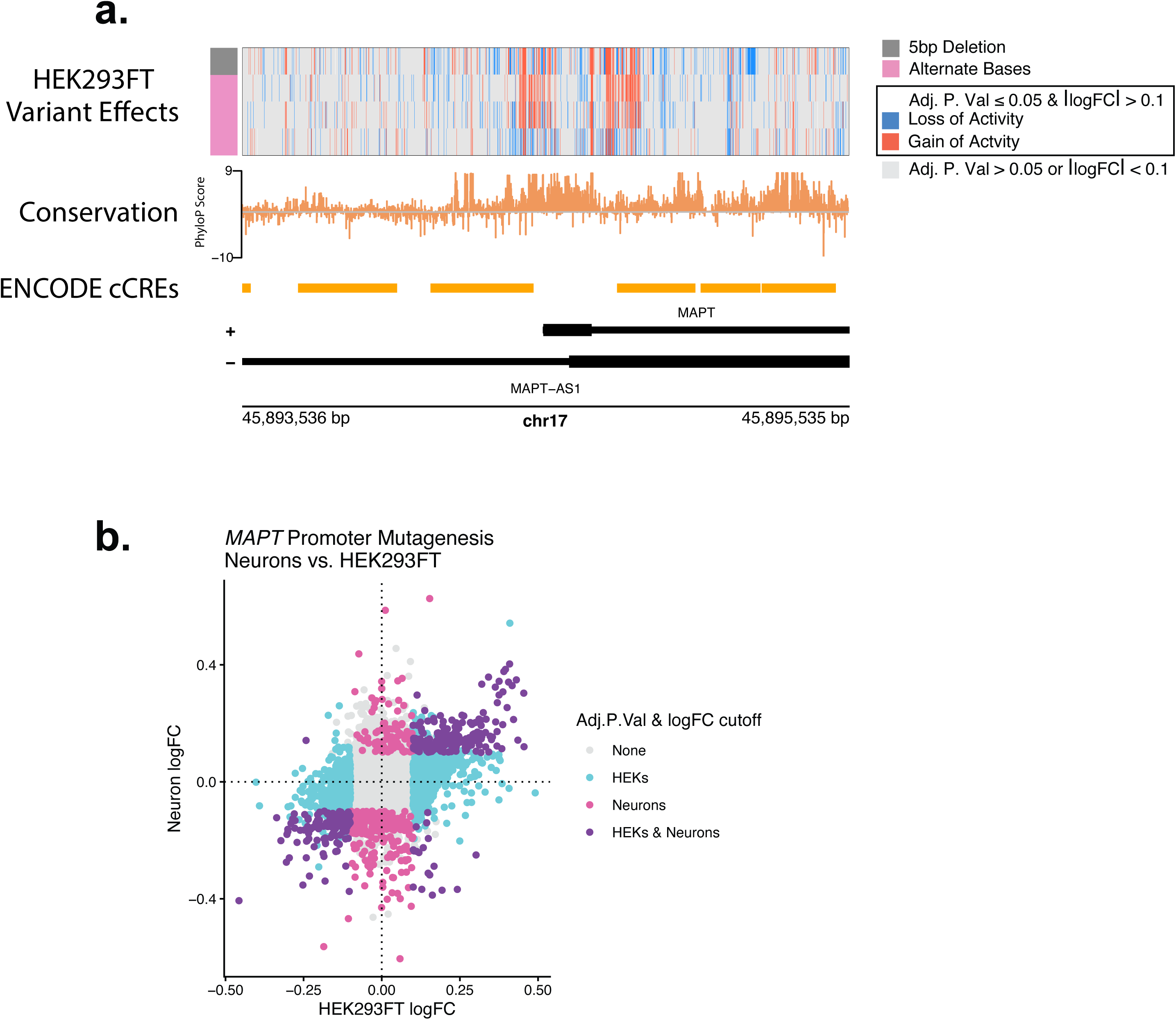
**a.** Gene track of the *MAPT* promoter saturation mutagenesis region showing variant effects observed in HEK293FT cells (pval.mad ≤ 0.05 and |logFC| > 0.1). The top row of the heatmap represents the centered 5 bp deletion sequences, and the bottom 3 rows represent each alternate base at that position. Red lines represent variants with a gain of activity and blue lines represent a loss of activity. **b.** Correlation plot of logFC of variant effects observed in Neurons and HEK293FT cells (pval.mad ≤ 0.05 and a |logFC| of at least 0.1). Blue dots represent variant effects in HEK293FT cells, pink represent variant effects in Neurons, and purple are variant effects in both cell types. (logFC Spearman’s ⍴ = 0.336, approximate p < 2.2×10^-16^)

**Figure s4.**
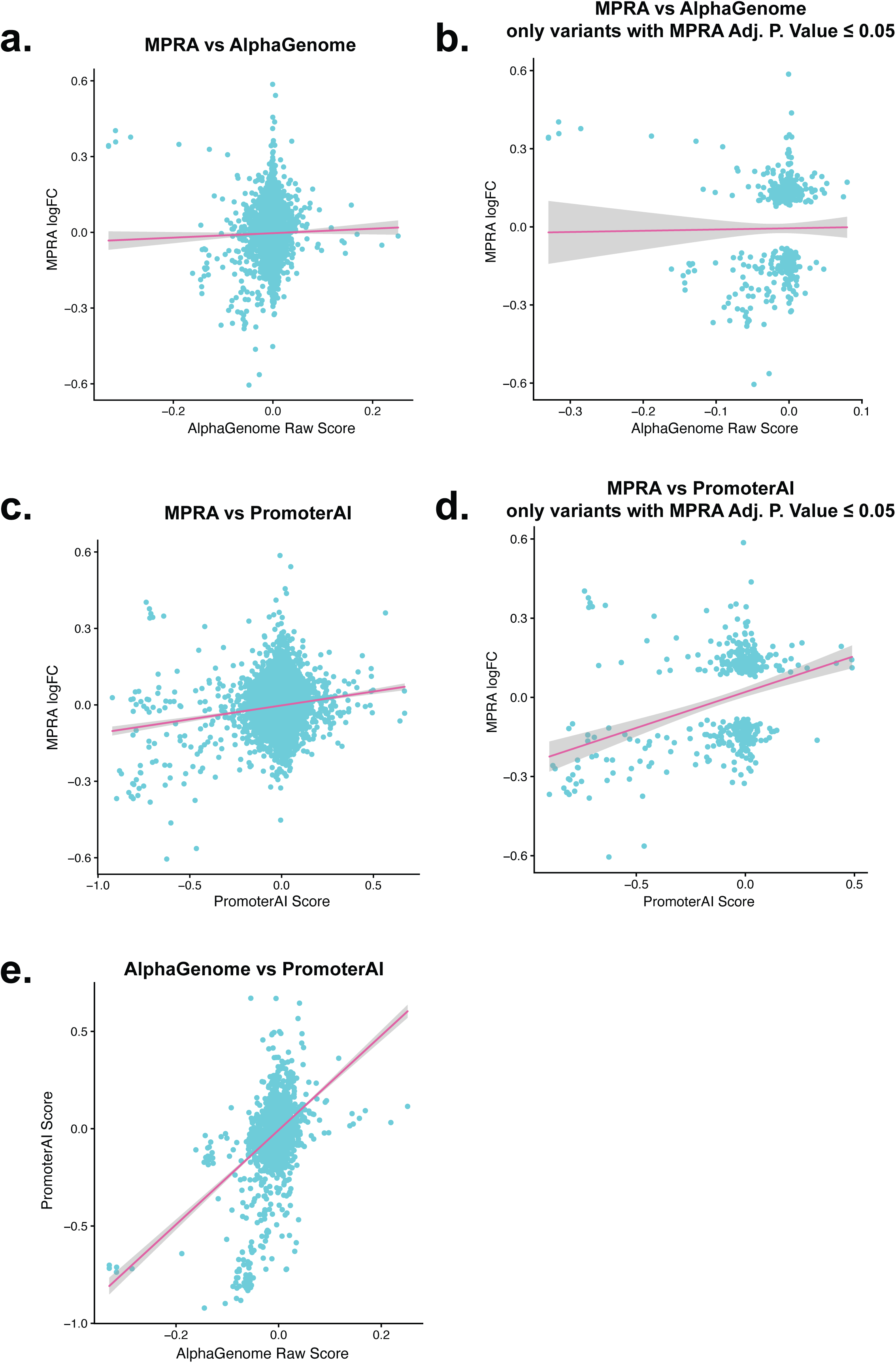
**a.** Correlation plots of AlphaGenome raw scores and MPRA logFC from the *MAPT* promoter mutagenesis. (Spearman’s ⍴ = 0.0277, approximate p = 0.0321) **b.** Same as a. except only variants with an adj. p. value ≤ 0.05 in the Neuron MPRA by bcalm are plotted. (Spearman’s ⍴ = 8.71×10^-9^, approximate p = 8.71×10^-4^) **c.** Correlation plots of PromoterAI scores and MPRA logFC from the *MAPT* promoter mutagenesis. (Spearman’s ⍴ = 0.482, approximate p = 1.89×10^-4^) **d.** Same as in c except only variants with an adj. p. value ≤ 0.05 in the Neuron MPRA by bcalm are plotted. (Spearman’s ⍴ = 0.2924, approximate p = 1.10×10^-9^) **e.** Correlation plot of AlphaGenome raw scores and PromoterAI scores for each SNV in the saturation mutagenesis MPRA (Spearman’s ⍴ = 0.153, approximate p < 2.2×10^-16^). For all plots, the best fit line is a linear regression of the data.

## Extended Acknowledgements

Data for this study were prepared, archived, and distributed by the National Institute on Aging Alzheimer’s Disease Data Storage Site (NIAGADS) at the University of Pennsylvania (U24-AG041689), funded by the National Institute on Aging.

The Alzheimer’s Disease Sequencing Project (ADSP) is comprised of two Alzheimer’s Disease (AD) genetics consortia and three National Human Genome Research Institute (NHGRI) funded Large Scale Sequencing and Analysis Centers (LSAC). The two AD genetics consortia are the Alzheimer’s Disease Genetics Consortium (ADGC) funded by NIA (U01 AG032984), and the Cohorts for Heart and Aging Research in Genomic Epidemiology (CHARGE) funded by NIA (R01 AG033193), the National Heart, Lung, and Blood Institute (NHLBI), other National Institute of Health (NIH) institutes and other foreign governmental and non-governmental organizations. The Discovery Phase analysis of sequence data is supported through UF1AG047133 (to Drs. Schellenberg, Farrer, Pericak-Vance, Mayeux, and Haines); U01AG049505 to Dr. Seshadri; U01AG049506 to Dr. Boerwinkle; U01AG049507 to Dr. Wijsman; and U01AG049508 to Dr. Goate and the Discovery Extension Phase analysis is supported through U01AG052411 to Dr. Goate, U01AG052410 to Dr. Pericak-Vance and U01 AG052409 to Drs. Seshadri and Fornage.

Sequencing for the Follow Up Study (FUS) is supported through U01AG057659 (to Drs. PericakVance, Mayeux, and Vardarajan) and U01AG062943 (to Drs. Pericak-Vance and Mayeux). Data generation and harmonization in the Follow-up Phase is supported by U54AG052427 (to Drs. Schellenberg and Wang). The FUS Phase analysis of sequence data is supported through U01AG058589 (to Drs. Destefano, Boerwinkle, De Jager, Fornage, Seshadri, and Wijsman), U01AG058654 (to Drs. Haines, Bush, Farrer, Martin, and Pericak-Vance), U01AG058635 (to Dr. Goate), RF1AG058066 (to Drs. Haines, Pericak-Vance, and Scott), RF1AG057519 (to Drs. Farrer and Jun), R01AG048927 (to Dr. Farrer), and RF1AG054074 (to Drs. Pericak-Vance and Beecham). The ADGC cohorts include: Adult Changes in Thought (ACT) (U01 AG006781, U19 AG066567), the Alzheimer’s Disease Research Centers (ADRC) (P30 AG062429, P30 AG066468, P30 AG062421, P30 AG066509, P30 AG066514, P30 AG066530, P30 AG066507, P30 AG066444, P30 AG066518, P30 AG066512, P30 AG066462, P30 AG072979, P30 AG072972, P30 AG072976, P30 AG072975, P30 AG072978, P30 AG072977, P30 AG066519, P30 AG062677, P30 AG079280, P30 AG062422, P30 AG066511, P30 AG072946, P30 AG062715, P30 AG072973, P30 AG066506, P30 AG066508, P30 AG066515, P30 AG072947, P30 AG072931, P30 AG066546, P20 AG068024, P20 AG068053, P20 AG068077, P20 AG068082, P30 AG072958, P30 AG072959), the Chicago Health and Aging Project (CHAP) (R01 AG11101, RC4 AG039085, K23 AG030944), Indiana Memory and Aging Study (IMAS) (R01 AG019771), Indianapolis Ibadan (R01 AG009956, P30 AG010133), the Memory and Aging Project (MAP) (R01 AG17917), Mayo Clinic (MAYO) (R01 AG032990, U01 AG046139, R01 NS080820, RF1 AG051504, P50 AG016574), Mayo Parkinson’s Disease controls (NS039764, NS071674, 5RC2HG005605), University of Miami (R01 AG027944, R01 AG028786, R01 AG019085, IIRG09133827, A2011048), the Multi-Institutional Research in Alzheimer’s Genetic Epidemiology Study (MIRAGE) (R01 AG09029, R01 AG025259), the National Centralized Repository for Alzheimer’s Disease and Related Dementias (NCRAD) (U24 AG021886), the National Institute on Aging Late Onset Alzheimer’s Disease Family Study (NIA-LOAD) (U24 AG056270), the Religious Orders Study (ROS) (P30 AG10161, R01 AG15819), the Texas Alzheimer’s Research and Care Consortium (TARCC) (funded by the Darrell K Royal Texas Alzheimer’s Initiative), Vanderbilt University/Case Western Reserve University (VAN/CWRU) (R01 AG019757, R01 AG021547, R01 AG027944, R01 AG028786, P01 NS026630, and Alzheimer’s Association), the Washington Heights-Inwood Columbia Aging Project (WHICAP) (RF1 AG054023), the University of Washington Families (VA Research Merit Grant, NIA: P50AG005136, R01AG041797, NINDS: R01NS069719), the Columbia University Hispanic Estudio Familiar de Influencia Genetica de Alzheimer (EFIGA) (RF1 AG015473), the University of Toronto (UT) (funded by Wellcome Trust, Medical Research Council, Canadian Institutes of Health Research), and Genetic Differences (GD) (R01 AG007584). The CHARGE cohorts are supported in part by National Heart, Lung, and Blood Institute (NHLBI) infrastructure grant HL105756 (Psaty), RC2HL102419 (Boerwinkle) and the neurology working group is supported by the National Institute on Aging (NIA) R01 grant AG033193.

The CHARGE cohorts participating in the ADSP include the following: Austrian Stroke Prevention Study (ASPS), ASPS-Family study, and the Prospective Dementia Registry-Austria (ASPS/PRODEM-Aus), the Atherosclerosis Risk in Communities (ARIC) Study, the Cardiovascular Health Study (CHS), the Erasmus Rucphen Family Study (ERF), the Framingham Heart Study (FHS), and the Rotterdam Study (RS). ASPS is funded by the Austrian Science Fond (FWF) grant number P20545-P05 and P13180 and the Medical University of Graz. The ASPS-Fam is funded by the Austrian Science Fund (FWF) project I904), the EU Joint Programme – Neurodegenerative Disease Research (JPND) in frame of the BRIDGET project (Austria, Ministry of Science) and the Medical University of Graz and the Steiermärkische Krankenanstalten Gesellschaft. PRODEM-Austria is supported by the Austrian Research Promotion agency (FFG) (Project No. 827462) and by the Austrian National Bank (Anniversary Fund, project 15435. ARIC research is carried out as a collaborative study supported by NHLBI contracts (HHSN268201100005C, HHSN268201100006C, HHSN268201100007C, HHSN268201100008C, HHSN268201100009C, HHSN268201100010C, HHSN268201100011C, and HHSN268201100012C). Neurocognitive data in ARIC is collected by U01 2U01HL096812, 2U01HL096814, 2U01HL096899, 2U01HL096902, 2U01HL096917 from the NIH (NHLBI, NINDS, NIA and NIDCD), and with previous brain MRI examinations funded by R01-HL70825 from the NHLBI. CHS research was supported by contracts HHSN268201200036C, HHSN268200800007C, N01HC55222, N01HC85079, N01HC85080, N01HC85081, N01HC85082, N01HC85083, N01HC85086, and grants U01HL080295 and U01HL130114 from the NHLBI with additional contribution from the National Institute of Neurological Disorders and Stroke (NINDS). Additional support was provided by R01AG023629, R01AG15928, and R01AG20098 from the NIA. FHS research is supported by NHLBI contracts N01-HC-25195 and HHSN268201500001I. This study was also supported by additional grants from the NIA (R01s AG054076, AG049607 and AG033040 and NINDS (R01 NS017950). The ERF study as a part of EUROSPAN (European Special Populations Research Network) was supported by European Commission FP6 STRP grant number 018947 (LSHG-CT-2006-01947) and also received funding from the European Community’s Seventh Framework Programme (FP7/2007-2013)/grant agreement HEALTH-F4- 2007-201413 by the European Commission under the programme “Quality of Life and Management of the Living Resources” of 5th Framework Programme (no. QLG2-CT-2002- 01254). High-throughput analysis of the ERF data was supported by a joint grant from the Netherlands Organization for Scientific Research and the Russian Foundation for Basic Research (NWO-RFBR 047.017.043). The Rotterdam Study is funded by Erasmus Medical Center and Erasmus University, Rotterdam, the Netherlands Organization for Health Research and Development (ZonMw), the Research Institute for Diseases in the Elderly (RIDE), the Ministry of Education, Culture and Science, the Ministry for Health, Welfare and Sports, the European Commission (DG XII), and the municipality of Rotterdam. Genetic data sets are also supported by the Netherlands Organization of Scientific Research NWO Investments (175.010.2005.011, 911-03-012), the Genetic Laboratory of the Department of Internal Medicine, Erasmus MC, the Research Institute for Diseases in the Elderly (014-93-015; RIDE2), and the Netherlands Genomics Initiative (NGI)/Netherlands Organization for Scientific Research (NWO) Netherlands Consortium for Healthy Aging (NCHA), project 050-060-810. All studies are grateful to their participants, faculty and staff. The content of these manuscripts is solely the responsibility of the authors and does not necessarily represent the official views of the National Institutes of Health or the U.S. Department of Health and Human Services.

The FUS cohorts include: the Alzheimer’s Disease Research Centers (ADRC) (P30 AG062429, P30 AG066468, P30 AG062421, P30 AG066509, P30 AG066514, P30 AG066530, P30 AG066507, P30 AG066444, P30 AG066518, P30 AG066512, P30 AG066462, P30 AG072979, P30 AG072972, P30 AG072976, P30 AG072975, P30 AG072978, P30 AG072977, P30 AG066519, P30 AG062677, P30 AG079280, P30 AG062422, P30 AG066511, P30 AG072946, P30 AG062715, P30 AG072973, P30 AG066506, P30 AG066508, P30 AG066515, P30 AG072947, P30 AG072931, P30 AG066546, P20 AG068024, P20 AG068053, P20 AG068077, P20 AG068082, P30 AG072958, P30 AG072959), Alzheimer’s Disease Neuroimaging Initiative (ADNI) (U19AG024904), Amish Protective Variant Study (RF1AG058066), Cache County Study (R01AG11380, R01AG031272, R01AG21136, RF1AG054052), Case Western Reserve University Brain Bank (CWRUBB) (P50AG008012), Case Western Reserve University Rapid Decline (CWRURD) (RF1AG058267, NU38CK000480), CubanAmerican Alzheimer’s Disease Initiative (CuAADI) (3U01AG052410), Estudio Familiar de Influencia Genetica en Alzheimer (EFIGA) (5R37AG015473, RF1AG015473, R56AG051876), Genetic and Environmental Risk Factors for Alzheimer Disease Among African Americans Study (GenerAAtions) (2R01AG09029, R01AG025259, 2R01AG048927), Gwangju Alzheimer and Related Dementias Study (GARD) (U01AG062602), Hillblom Aging Network (2014-A-004-NET, R01AG032289, R01AG048234), Hussman Institute for Human Genomics Brain Bank (HIHGBB) (R01AG027944, Alzheimer’s Association “Identification of Rare Variants in Alzheimer Disease”), Ibadan Study of Aging (IBADAN) (5R01AG009956), Longevity Genes Project (LGP) and LonGenity (R01AG042188, R01AG044829, R01AG046949, R01AG057909, R01AG061155, P30AG038072), Mexican Health and Aging Study (MHAS) (R01AG018016), Multi-Institutional Research in Alzheimer’s Genetic Epidemiology (MIRAGE) (2R01AG09029, R01AG025259, 2R01AG048927), Northern Manhattan Study (NOMAS) (R01NS29993), Peru Alzheimer’s Disease Initiative (PeADI) (RF1AG054074), Puerto Rican 1066 (PR1066) (Wellcome Trust (GR066133/GR080002), European Research Council (340755)), Puerto Rican Alzheimer Disease Initiative (PRADI) (RF1AG054074), Reasons for Geographic and Racial Differences in Stroke (REGARDS) (U01NS041588), Research in African American Alzheimer Disease Initiative (REAAADI) (U01AG052410), the Religious Orders Study (ROS) (P30 AG10161, P30 AG72975, R01 AG15819, R01 AG42210), the RUSH Memory and Aging Project (MAP) (R01 AG017917, R01 AG42210Stanford Extreme Phenotypes in AD (R01AG060747), University of Miami Brain Endowment Bank (MBB), University of Miami/Case Western/North Carolina A&T African American (UM/CASE/NCAT) (U01AG052410, R01AG028786), Wisconsin Registry for Alzheimer’s Prevention (WRAP) (R01AG027161 and R01AG054047), Mexico-Southern California Autosomal Dominant Alzheimer’s Disease Consortium (R01AG069013), Center for Cognitive Neuroscience and Aging (R01AG047649), and the A4 Study (R01AG063689, U19AG010483 and U24AG057437).

The four LSACs are: the Human Genome Sequencing Center at the Baylor College of Medicine (U54 HG003273), the Broad Institute Genome Center (U54HG003067), The American Genome Center at the Uniformed Services University of the Health Sciences (U01AG057659), and the Washington University Genome Institute (U54HG003079). Genotyping and sequencing for the ADSP FUS is also conducted at John P. Hussman Institute for Human Genomics (HIHG) Center for Genome Technology (CGT).

Biological samples and associated phenotypic data used in primary data analyses were stored at Study Investigators institutions, and at the National Centralized Repository for Alzheimer’s Disease and Related Dementias (NCRAD, U24AG021886) at Indiana University funded by NIA. Associated Phenotypic Data used in primary and secondary data analyses were provided by Study Investigators, the NIA funded Alzheimer’s Disease Centers (ADCs), and the National Alzheimer’s Coordinating Center (NACC, U24AG072122) and the National Institute on Aging Genetics of Alzheimer’s Disease Data Storage Site (NIAGADS, U24AG041689) at the University of Pennsylvania, funded by NIA. Harmonized phenotypes were provided by the ADSP Phenotype Harmonization Consortium (ADSP-PHC), funded by NIA (U24 AG074855, U01 AG068057 and R01 AG059716) and Ultrascale Machine Learning to Empower Discovery in Alzheimer’s Disease Biobanks (AI4AD, U01 AG068057). This research was supported in part by the Intramural Research Program of the National Institutes of health, National Library of Medicine. Contributors to the Genetic Analysis Data included Study Investigators on projects that were individually funded by NIA, and other NIH institutes, and by private U.S. organizations, or foreign governmental or nongovernmental organizations.

The ADSP Phenotype Harmonization Consortium (ADSP-PHC) is funded by NIA (U24 AG074855, U01 AG068057 and R01 AG059716). The harmonized cohorts within the ADSP-PHC include: the Anti-Amyloid Treatment in Asymptomatic Alzheimer’s study (A4 Study), a secondary prevention trial in preclinical Alzheimer’s disease, aiming to slow cognitive decline associated with brain amyloid accumulation in clinically normal older individuals. The A4 Study is funded by a public-private-philanthropic partnership, including funding from the National Institutes of Health-National Institute on Aging, Eli Lilly and Company, Alzheimer’s Association, Accelerating Medicines Partnership, GHR Foundation, an anonymous foundation and additional private donors, with in-kind support from Avid and Cogstate. The companion observational Longitudinal Evaluation of Amyloid Risk and Neurodegeneration (LEARN) Study is funded by the Alzheimer’s Association and GHR Foundation. The A4 and LEARN Studies are led by Dr. Reisa Sperling at Brigham and Women’s Hospital, Harvard Medical School and Dr. Paul Aisen at the Alzheimer’s Therapeutic Research Institute (ATRI), University of Southern California. The A4 and LEARN Studies are coordinated by ATRI at the University of Southern California, and the data are made available through the Laboratory for Neuro Imaging at the University of Southern California. The participants screening for the A4 Study provided permission to share their de-identified data in order to advance the quest to find a successful treatment for Alzheimer’s disease. We would like to acknowledge the dedication of all the participants, the site personnel, and all of the partnership team members who continue to make the A4 and LEARN Studies possible. The complete A4 Study Team list is available on: a4study.org/a4-study-team.; the Adult Changes in Thought study (ACT), U01 AG006781, U19 AG066567; Alzheimer’s Disease Neuroimaging Initiative (ADNI): Data collection and sharing for this project was funded by the Alzheimer’s Disease Neuroimaging Initiative (ADNI) (National Institutes of Health Grant U01 AG024904) and DOD ADNI (Department of Defense award number W81XWH-12-2-0012). ADNI is funded by the National Institute on Aging, the National Institute of Biomedical Imaging and Bioengineering, and through generous contributions from the following: AbbVie, Alzheimer’s Association; Alzheimer’s Drug Discovery Foundation; Araclon Biotech; BioClinica, Inc.; Biogen; Bristol-Myers Squibb Company; CereSpir, Inc.; Cogstate; Eisai Inc.; Elan Pharmaceuticals, Inc.; Eli Lilly and Company; EuroImmun; F. Hoffmann-La Roche Ltd and its affiliated company Genentech, Inc.; Fujirebio; GE Healthcare; IXICO Ltd.;Janssen Alzheimer Immunotherapy Research & Development, LLC.; Johnson & Johnson Pharmaceutical Research & Development LLC.; Lumosity; Lundbeck; Merck & Co., Inc.;Meso Scale Diagnostics, LLC.; NeuroRx Research; Neurotrack Technologies; Novartis Pharmaceuticals Corporation; Pfizer Inc.; Piramal Imaging; Servier; Takeda Pharmaceutical Company; and Transition Therapeutics. The Canadian Institutes of Health Research is providing funds to support ADNI clinical sites in Canada. Private sector contributions are facilitated by the Foundation for the National Institutes of Health (www.fnih.org). The grantee organization is the Northern California Institute for Research and Education, and the study is coordinated by the Alzheimer’s Therapeutic Research Institute at the University of Southern California. ADNI data are disseminated by the Laboratory for Neuro Imaging at the University of Southern California; Estudio Familiar de Influencia Genetica en Alzheimer (EFIGA): 5R37AG015473, RF1AG015473, R56AG051876; the Health & Aging Brain Study – Health Disparities (HABS-HD), supported by the National Institute on Aging of the National Institutes of Health under Award Numbers R01AG054073, R01AG058533, R01AG070862, P41EB015922, and U19AG078109; the Korean Brain Aging Study for the Early Diagnosis and Prediction of Alzheimer’s disease (KBASE), which was supported by a grant from Ministry of Science, ICT and Future Planning (Grant No: NRF-2014M3C7A1046042); Memory & Aging Project at Knight Alzheimer’s Disease Research Center (MAP at Knight ADRC): The Memory and Aging Project at the Knight-ADRC (Knight-ADRC). This work was supported by the National Institutes of Health (NIH) grants R01AG064614, R01AG044546, RF1AG053303, RF1AG058501, U01AG058922 and R01AG064877 to Carlos Cruchaga. The recruitment and clinical characterization of research participants at Washington University was supported by NIH grants P30AG066444, P01AG03991, and P01AG026276. Data collection and sharing for this project was supported by NIH grants RF1AG054080, P30AG066462, R01AG064614 and U01AG052410. We thank the contributors who collected samples used in this study, as well as patients and their families, whose help and participation made this work possible. This work was supported by access to equipment made possible by the Hope Center for Neurological Disorders, the Neurogenomics and Informatics Center (NGI: https://neurogenomics.wustl.edu/) and the Departments of Neurology and Psychiatry at Washington University School of Medicine; National Alzheimer’s Coordinating Center (NACC): The NACC database is funded by NIA/NIH Grant U24 AG072122. SCAN is a multi-institutional project that was funded as a U24 grant (AG067418) by the National Institute on Aging in May 2020. Data collected by SCAN and shared by NACC are contributed by the NIA-funded ADRCs as follows: P30 AG062429 (PI James Brewer, MD, PhD), P30 AG066468 (PI Oscar Lopez, MD), P30 AG062421 (PI Bradley Hyman, MD, PhD), P30 AG066509 (PI Thomas Grabowski, MD), P30 AG066514 (PI Mary Sano, PhD), P30 AG066530 (PI Helena Chui, MD), P30 AG066507 (PI Marilyn Albert, PhD), P30 AG066444 (PI John Morris, MD), P30 AG066518 (PI Jeffrey Kaye, MD), P30 AG066512 (PI Thomas Wisniewski, MD), P30 AG066462 (PI Scott Small, MD), P30 AG072979 (PI David Wolk, MD), P30 AG072972 (PI Charles DeCarli, MD), P30 AG072976 (PI Andrew Saykin, PsyD), P30 AG072975 (PI David Bennett, MD), P30 AG072978 (PI Neil Kowall, MD), P30 AG072977 (PI Robert Vassar, PhD), P30 AG066519 (PI Frank LaFerla, PhD), P30 AG062677 (PI Ronald Petersen, MD, PhD), P30 AG079280 (PI Eric Reiman, MD), P30 AG062422 (PI Gil Rabinovici, MD), P30 AG066511 (PI Allan Levey, MD, PhD), P30 AG072946 (PI Linda Van Eldik, PhD), P30 AG062715 (PI Sanjay Asthana, MD, FRCP), P30 AG072973 (PI Russell Swerdlow, MD), P30 AG066506 (PI Todd Golde, MD, PhD), P30 AG066508 (PI Stephen Strittmatter, MD, PhD), P30 AG066515 (PI Victor Henderson, MD, MS), P30 AG072947 (PI Suzanne Craft, PhD), P30 AG072931 (PI Henry Paulson, MD, PhD), P30 AG066546 (PI Sudha Seshadri, MD), P20 AG068024 (PI Erik Roberson, MD, PhD), P20 AG068053 (PI Justin Miller, PhD), P20 AG068077 (PI Gary Rosenberg, MD), P20 AG068082 (PI Angela Jefferson, PhD), P30 AG072958 (PI Heather Whitson, MD), P30 AG072959 (PI James Leverenz, MD); National Institute on Aging Alzheimer’s Disease Family Based Study (NIA-AD FBS): U24 AG056270; Religious Orders Study (ROS): P30AG10161,R01AG15819, R01AG42210; Memory and Aging Project (MAP - Rush): R01AG017917, R01AG42210; Minority Aging Research Study (MARS): R01AG22018, R01AG42210; the Texas Alzheimer’s Research and Care Consortium (TARCC), funded by the Darrell K Royal Texas Alzheimer’s Initiative, directed by the Texas Council on Alzheimer’s Disease and Related Disorders; Washington Heights/Inwood Columbia Aging Project (WHICAP): RF1 AG054023;and Wisconsin Registry for Alzheimer’s Prevention (WRAP): R01AG027161 and R01AG054047. Additional acknowledgments include the National Institute on Aging Genetics of Alzheimer’s Disease Data Storage Site (NIAGADS, U24AG041689) at the University of Pennsylvania, funded by NIA.

## BioRender Citations

Figure 1a, LentiMPRA overview: Created in BioRender. Gordon, E. (2026) https://BioRender.com/h153oek

Figure 1b, BAC MPRA: Created in BioRender. Gordon, E. (2026) https://BioRender.com/qjavnet

Figure 1b, Oligo MPRA: Created in BioRender. Gordon, E. (2026) https://BioRender.com/nrlydup

Figure 4a, Saturation Mutagenesis Design: Created in BioRender. Gordon, E. (2026) https://BioRender.com/7fyfi82

